# Polyubiquitinated PCNA promotes alternative lengthening of telomeres by inducing break-induced replication

**DOI:** 10.1101/2023.07.13.548953

**Authors:** Sangin Kim, Nalae Kang, Jae Sun Ra, Su Hyung Park, Kyungjae Myung, Kyoo-young Lee

## Abstract

Replication stresses are the major source of break-induced replication (BIR). Here, we show that in alternative lengthening of telomeres (ALT) cells, replication stress-induced polyubiquitinated PCNA (polyUb-PCNA) triggers BIR at telomeres and the common fragile site (CFS). Consistently, depleting RAD18, a PCNA ubiquitinating enzyme, reduces the occurrence of ALT-associated PML bodies (APBs) and mitotic DNA synthesis at telomeres and CFS, both of which are mediated by BIR. In contrast, inhibiting USP1, an Ub-PCNA deubiquitinating enzyme, results in an increase in the above phenotypes in a RAD18- and UBC13 (the PCNA polyubiquitinating enzyme)-dependent manner. Furthermore, deficiency of ATAD5, which facilitates USP1 activity and unloads PCNAs, augments recombination-associated phenotypes. Mechanistically, telomeric polyUb-PCNA accumulates SLX4, a nuclease scaffold, at telomeres through its ubiquitin-binding domain and increases telomere damage. Consistently, APB increase induced by Ub-PCNA depends on SLX4 and SLX4-associated nucleases. Taken together, our results identified the polyUb-PCNA-SLX4 axis as a trigger for directing BIR.

## Introduction

Eukaryotic somatic cells undergo telomere shortening due to loss of telomerase (TEL) activity (De Vitis et al., 2018; Shay, 2016). The majority of human cancers maintain telomere length by restoring TEL activity (Kim et al., 1994). Furthermore, 10∼15% of human cancers adopt a recombination-dependent alternative lengthening of telomeres (ALT) pathway instead of TEL activity (Bryan et al., 1997; Cesare and Reddel, 2010). ALT involves conservative telomeric DNA synthesis occurs via a break-induced replication (BIR) that depends on POLD3 and RAD52 recombinase (Anand et al., 2013; Costantino et al., 2014; Malkova and Ira, 2013; Min et al., 2017; Roumelioti et al., 2016; Zhang et al., 2019; Zhang and Zou, 2020). ALT+ cancer cells feature ALT-associated promyelocytic leukemia (PML) body (APB), containing telomere clusters and proteins for recombination and DNA synthesis (Draskovic et al., 2009; Min et al., 2017; Yeager et al., 1999; Zhang et al., 2019; Zhang and Zou, 2020).

Replication stress affects difficult-to-replicate regions like common fragile sites (CFSs) or telomeres (Bhowmick and Hickson, 2017; Maffia et al., 2020; Petermann et al., 2022). In CFSs, when cells enter mitosis with incomplete DNA replication under replication stress, RAD52-dependent BIR-like mitotic DNA synthesis (MiDAS) occurs (Bhowmick et al., 2016). Telomeric replication stress leads to telomere clustering and also MiDAS at telomeres (Min et al., 2017; Ozer et al., 2018). Telomeric R-loop and G-quadruplex (G4) levels are higher in ALT+ cells compared to TEL+ cancer cells (Yang et al., 2021a). Despite evidence that telomeric replication stresses trigger ALT activity (Amato et al., 2020; Arora et al., 2014; Pan et al., 2019; Pan et al., 2017; Silva et al., 2019; Vohhodina et al., 2021; Yang et al., 2021a), it is unclear how replication stress results in the telomere breakage required to initiate the BIR-associated ALT process.

The eukaryotic sliding clamp proliferating cell nuclear antigen (PCNA), which is loaded onto DNA by the pentameric replication factor C (RFC) complex, is critical for DNA replication and repair (Moldovan et al., 2007). It undergoes monoubiquitination by the RAD6-RAD18 complex in response to replication stress, recruiting translesion synthesis (TLS) polymerases for error-prone DNA lesion bypass (Hoege et al., 2002). PCNA can also undergo lysine 63 (K63)-linked polyubiquitination by the UBC13/MMS2 ubiquitin-conjugating dimer and yeast Rad5 ubiquitin ligase homologues, promoting error-free DNA damage tolerance through template switching (Hoege et al., 2002; Motegi et al., 2008; Unk et al., 2008). ATAD5 (human ortholog of yeast Elg1)-RFC-like-complex (RLC) unloads PCNA and Ub-PCNA and facilitates PCNA deubiquitination by recruiting the ubiquitin-specific protease 1 (USP1)/USP1-associated factor (UAF1) complex (Kang et al., 2019; Kubota et al., 2013; Lee et al., 2013; Lee et al., 2010). ATAD5 is important for faithful DNA replication and repair and for maintaining genomic stability by preventing PCNA/Ub-PCNA accumulation on chromatin (Gali et al., 2018; Kim et al., 2020; Lee et al., 2013; Lee and Park, 2020; Lee et al., 2010; Park et al., 2019; Park et al., 2021).

Elg1 loss results in telomere elongation (Banerjee and Myung, 2004; Smolikov et al., 2004), and a role for PCNA or Ub-PCNA in promoting telomere elongation has been suggested in e*lg1Δ* mutants (Johnson et al., 2016). PCNA ubiquitination is implicated in BIR, as observed in yeast and recent studies on MiDAS and break-induced telomere synthesis (Lydeard et al., 2010; Wu et al., 2023; Zhang et al., 2023). However, the specific involvement of Ub-PCNA in the BIR-associated ALT process and the role of polyUb-PCNA in ALT activity remain unclear. Here, we investigated the effects of modulating ATAD5-USP1 activity specifically in the G2 phase of ALT+ cancer cells to determine how Ub-PCNA contributes to the BIR-associated ALT process.

## Results

### Ub-PCNA localization at APBs increases in ATAD5-depleted cells in the G2 phase

PCNA and RAD18, a PCNA ubiquitinating enzyme, localizes at APBs (Garcia-Exposito et al., 2016; Zhang et al., 2021), and ATAD5 participates in PCNA unloading and deubiquitination at replication forks or DNA damage sites (Fig. 1a). We therefore examined the effects of ATAD5 depletion on PCNA and Ub-PCNA levels at telomeres in the G2 phase using ALT+ U2OS cancer cell line with endogenous ATAD5 protein fused to an auxin-inducible degron (AID) (Extended Data Fig. 1a) (Park et al., 2021). We found that both PCNA and Ub-PCNA localized at APBs and that the level of localization was increased in ATAD5-depleted U2OS cells in the G2 phase (Fig. 1b-e).

**Fig. 1.**
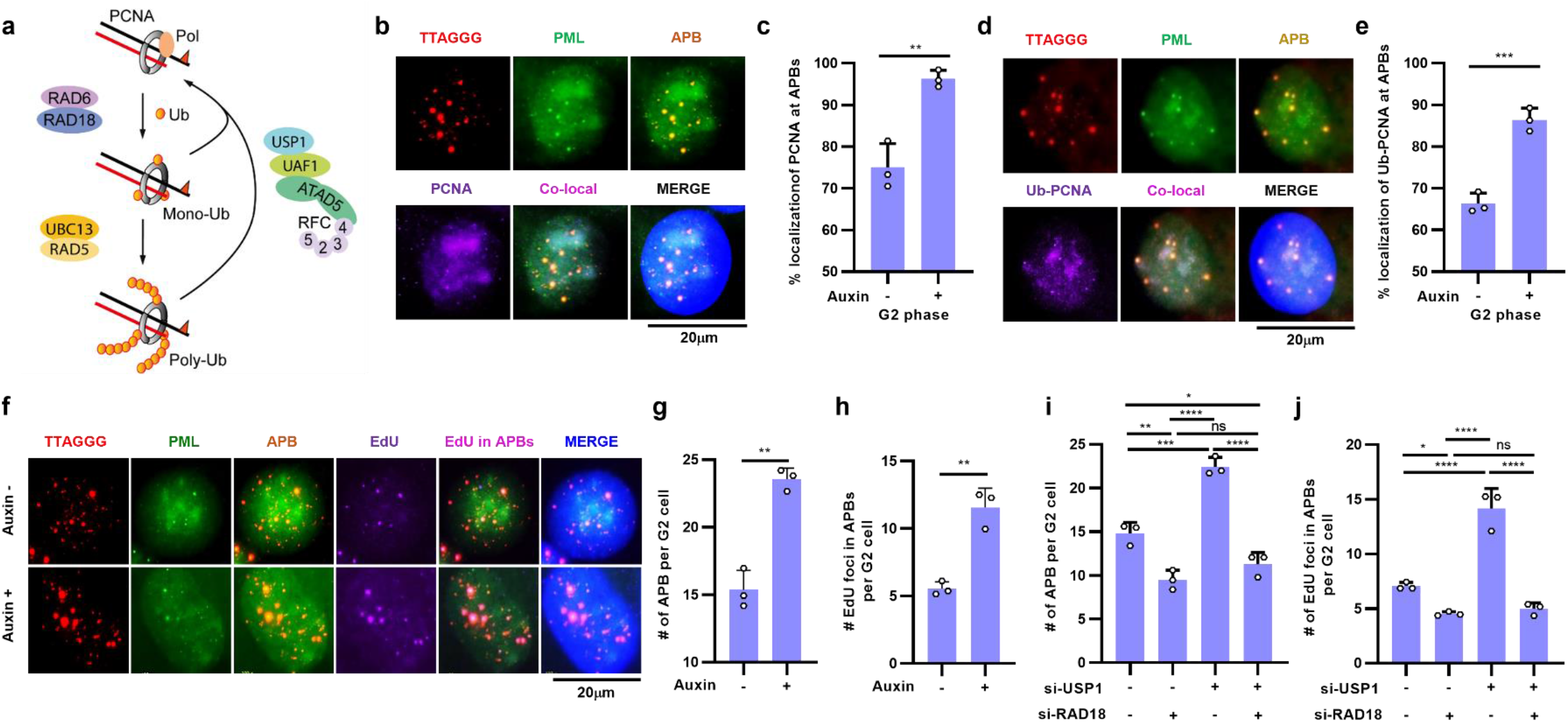
Ub-PCNA increases APBs with DNA synthesis in the G2 phase of ALT+ cancer cells. (a) A diagram of PCNA ubiquitination and de-ubiquitination. (b-e) U2OS-ATAD5^AID^ cells were treated with auxin for 48 h and fixed at the G2 phase for immunostaining with telomere FISH. (b, d) Representative images of localization of PCNA (b) or Ub-PCNA (d) at APBs. (c, e) The percent localization of PCNA (c) or Ub-PCNA (e) at APBs per G2 phase cell was quantified. (f-h) U2OS-ATAD5^AID^ cells were treated with auxin for 48 h and fixed at the G2 phase for an ATSA assay. (i, j) After siRNA transfection, G2-synchronized U2OS cells were subjected to an ATSA assay. (g-j) Quantification ATSA results were displayed. Data in all graphs represent the mean ± s.d. of at least three independent experiments. Statistical analysis: two-tailed unpaired Student’s *t*-test (c, e, g, h); One-way ANOVA (i, j).

### Ub-PCNA increases APBs with DNA synthesis in the G2 phase

Telomeric DNA synthesis can be analyzed using the ALT telomere DNA synthesis in APBs (ATSA) assay that detects intact telomere extension in ALT+ cells in the G2 phase (Zhang et al., 2019). As previously reported, 5′-ethynyl-2′-deoxyuridine (EdU) foci were detected in APBs of most U2OS cells in the G2 phase (Fig. 1f and Extended Data Fig. 1b, c). ATAD5 depletion increased the number of total APBs and EdU^+^ APBs (Fig. 1f-h). Consistent with the cooperative role of ATAD5 and USP1 in PCNA deubiquitination (Fig. 1a) (Lee et al., 2010), USP1 depletion also increased the number of total APBs and EdU^+^ APBs, which was reduced when RAD18 was co-depleted (Fig. 1i, j and Extended Data Fig. 1d). We found that RAD18 depletion alone decreased the number of total APBs and EdU^+^ APBs (Fig. 1i, j), which is in contrast to the increase in APB number by RAD18 depletion in asynchronous U2OS cells (Garcia-Exposito et al., 2016), suggesting that RAD18 depletion may have a differential effect on BIR-associated ALT depending on the cell cycle. Collectively, these results suggest that APB formation and telomeric DNA synthesis are restricted by PCNA deubiquitination.

### Ub-PCNA increases mitotic DNA synthesis events in ALT+ cancer cells

ALT-mediated telomeric DNA synthesis occurs via a BIR-like process (Dilley et al., 2016; Roumelioti et al., 2016; Zhang et al., 2019). Therefore, we investigated BIR-mediated MiDAS at telomeres and CFSs in the three ALT+ cell lines, U2OS, GM0637 and SK-LU-1 (Fig. 2a and Extended Data Fig. 2a-d) (Bhowmick et al., 2016; Min et al., 2017; Minocherhomji et al., 2015; Ozer et al., 2018). ATAD5 depletion increased MiDAS events at both telomeres and CFSs in U2OS cells under replication stress (Fig. 2b, c). Similar increases in MiDAS events at telomeres were observed in GM0637 and SK-LU-1 cells upon ATAD5 depletion (Fig. 2d, e). Co-depletion of POLD3 or RAD52 with ATAD5 restored MiDAS events to baseline levels in U2OS cells (Fig. 2f, g and Extended Data Fig. 2e, f). The dependence of this process on RAD52 was confirmed using *RAD52^−^*^/^*^−^* U2OS cells in which RAD52 expression can be induced by doxycycline (Zhang et al., 2019) (Fig. 2h and Extended Data Fig. 2g). Interestingly, ATAD5 depletion did not increase MiDAS at CFSs or telomeres in TEL+ HeLa_long telomere (HeLa_LT) cells (Fig. 2i, j and Extended Data Fig. 2h).

**Fig 2.**
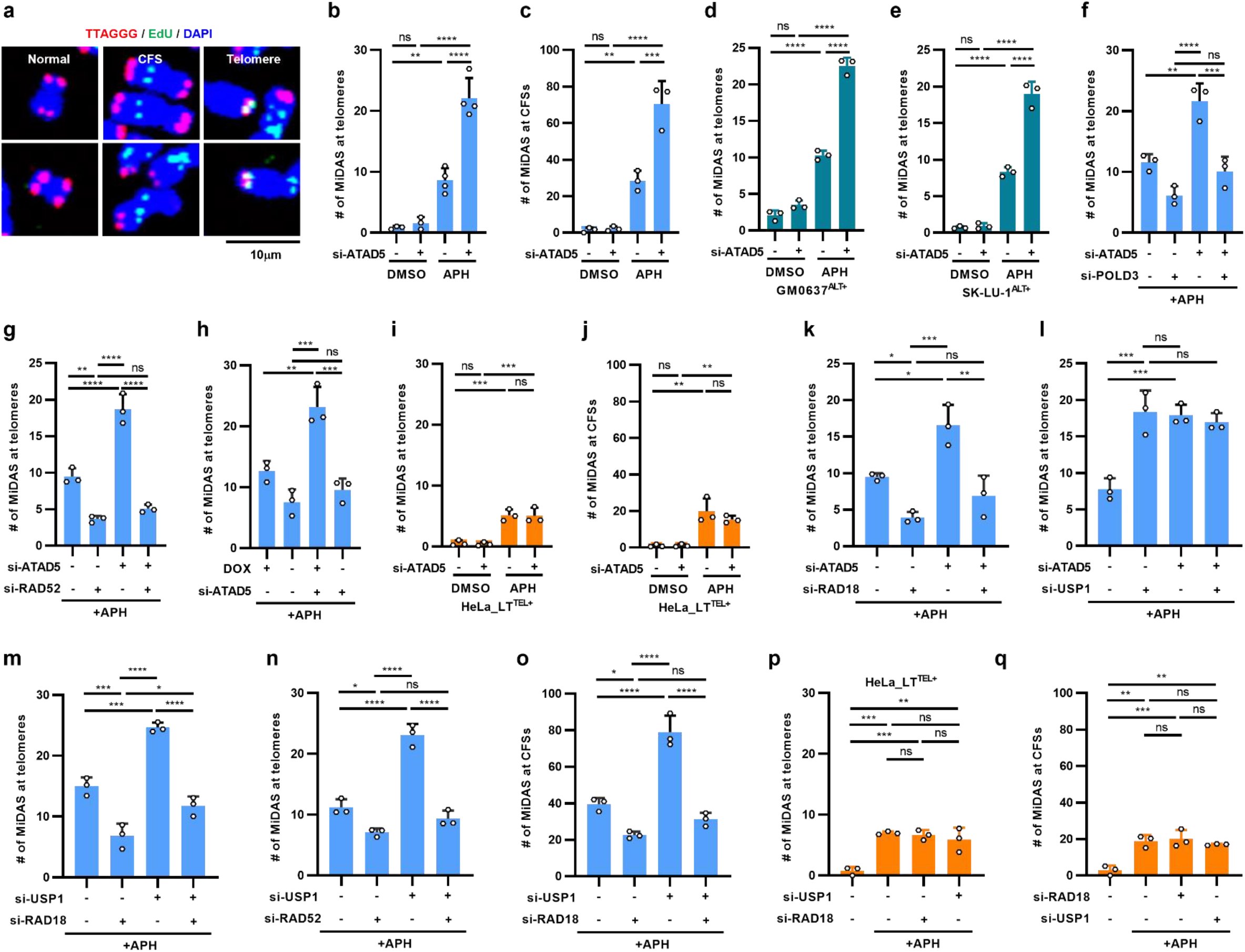
Ub-PCNA increases mitotic DNA synthesis events in ALT+ cancer cells. (a) Representative images of MiDAS after APH treatment. (a-q) After transfection, cells were treated with DMSO or 0.4 μM aphidicolin (APH) for 16 h and subjected to a MiDAS assay. (b-q) The number of MiDAS at telomeres or CFS regions was quantified. (h) RAD52 expression was induced by doxycycline (DOX) for 48 h. Cell lines: U2OS (b, c, f-h, k-o); GM0637 (d); SK-LU-1 (e); *RAD52*^−/−^ U2OS (h); and HeLa_LT (i, j, p, q). Data in all graphs represent the mean ± s.d. of at least three independent experiments. Statistical analysis: One-way ANOVA (b-q).

Consistent with the ATSA results, RAD18 depletion reduced the telomeric MiDAS induced by ATAD5 depletion in U2OS cells (Fig. 2k and Extended Data Fig. 2i). In contrast, USP1 depletion increased telomeric MiDAS to the level caused by ATAD5 single depletion, and this effect was not further increased by ATAD5 co-depletion (Fig. 2l and Extended Data Fig. 2j), but was restored to basal levels when either RAD18 or RAD52 was co-depleted (Fig. 2m, n and Extended Data Fig. 2k). MiDAS at CFSs was also similarly regulated by USP1 or RAD18 in U2OS cells (Fig. 2o). In contrast, MiDAS levels at telomeres and CFSs in HeLa_LT cells were not affected by USP1 or RAD18 depletion (Fig. 2p, q and Extended Data Fig. 2l). Considering that MiDAS in ALT+ and TEL+ cells share nearly the same BIR process (Minocherhomji et al., 2015; Ozer et al., 2018), these results collectively suggest that unprocessed Ub-PCNA increases BIR events only in ALT+ cancer cells.

### Ub-PCNA increases break-induced telomere synthesis

The telomeric repeat binding factor 1 (TRF1)-FokI endonuclease-induced telomere synthesis occurs in both ALT+ and TEL+ cells and the synthesis depends on PCNA and POLD3 (Fig. 3a) (Cho et al., 2014; Dilley et al., 2016). Upon induction of nuclear localization of the TRF1-FokI protein by tamoxifen treatment in the G2 phase, PCNA and EdU foci were colocalized with TRF1-FokI foci (Extended Data Fig. 3a), as reported (Dilley et al., 2016). Ub-PCNA also colocalized with TRF1-FokI and EdU foci upon inducing telomere breaks (Fig. 3b), which was significantly reduced by RAD18 depletion (Fig. 3c, d). Residual localization of Ub-PCNA at telomeric breaks might be explained by siRNA efficiency but not by RNF168-mediated PCNA ubiquitination (Yang et al., 2021b) (Extended Data Fig. 3b, c). We found that EdU signal at TRF1-FokI foci, which indicates break-induced telomere synthesis, was reduced by RAD18 depletion and increased by USP1 depletion (Fig. 3e, f).

**Fig 3.**
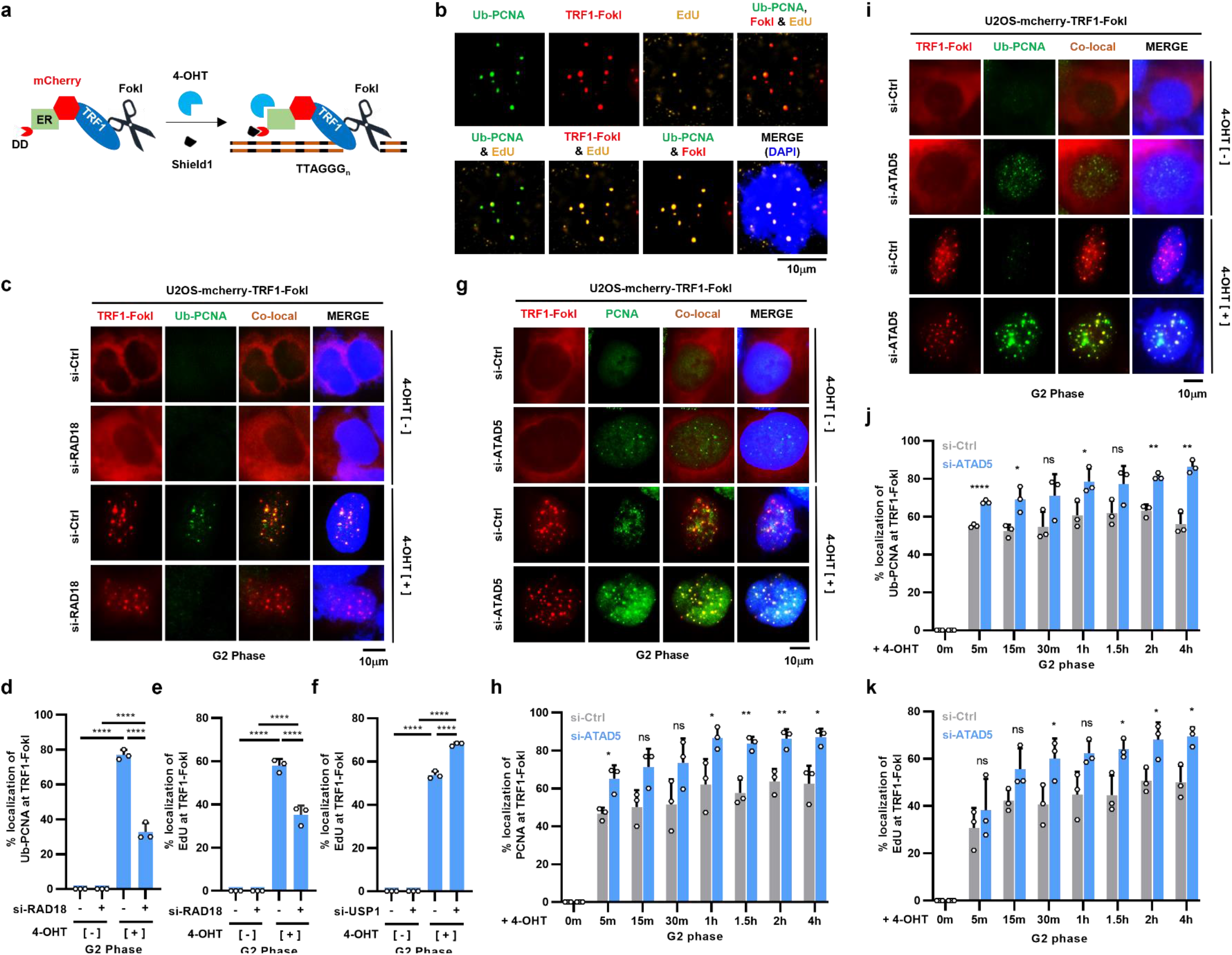
Ub-PCNA increases break-induced telomere synthesis. (a) A diagram of the TRF1-FokI system. (b-k) After siRNA transfection, U2OS-mCherry-TRF1-FokI cells were enriched in the G2 phase and treated with tamoxifen and Shield1 for 2 h (b-f, g, i) or indicated times (h, j, k) before fixation for immunostaining. (b) Representative immunostaining images of Ub-PCNA, EdU, and TRF1-FokI co-localization after telomeric break induction. (c, g, i) Representative images showing localization of Ub-PCNA (c, i) or PCNA (g) at TRF1-FokI foci. (d-f, h, j, k) The percent localization of Ub-PCNA (d, j), EdU (e, f, k) or PCNA (h) at TRF1-FokI foci was quantified. Data in all graphs represent the mean ± s.d. of at least three independent experiments. Statistical analysis: One-way ANOVA (d-f); two-tailed unpaired Student’s *t*-test (h, j, k).

PCNA and Ub-PCNA localization at TRF1-FokI foci was consistently increased in the G2 phase by ATAD5 depletion (Fig. 3g-j and Extended Data Fig. 3d-f). PCNA or Ub-PCNA signal was observed at TRF1-FokI foci starting at 5 min after break induction, and colocalization events were more frequent at all time points in ATAD5-depleted cells (Fig. 3h, j and Extended Data Fig. 3e, f). Similarly, EdU signal at TRF1-FokI foci was observed beginning at 5 min after break induction (Fig. 3k), suggesting a rapid onset of telomere synthesis. ATAD5 depletion resulted in an increase in telomere synthesis events at all time points (Fig. 3k). Taken together, these results suggest that PCNA ubiquitination contributes to break-induced telomere synthesis.

### ATAD5 deficiency extends telomere length in ALT+ cancer cells

ATAD5 depletion increased APBs with DNA synthesis and break-induced telomere synthesis, and e*lg1Δ* mutants exhibit telomere elongation (Banerjee and Myung, 2004; Smolikov et al., 2004). We therefore investigated the effect of long-term ATAD5 depletion on telomere length using ATAD5^AID^ cells (Extended Data Fig. 4a). We found that telomere lengthening started to appear in U2OS-ATAD5^AID^ cells treated with auxin for just 10 days (Fig. 4a). However, auxin treatment even for 40 days did not change the telomere length in HeLa-ATAD5^AID^ cells (Fig. 4b). In addition, *ATAD5^−^*^/^*^−^* U2OS and other ALT+ *ATAD5^−^*^/^*^−^* SK-LU-1 clones, but not TEL+ *ATAD5^−^*^/^*^−^* HeLa_LT clones, showed telomere lengthening (Fig. 4c and Extended Data Fig. 4b). In addition, we examined telomeric DNA extension using a single-molecule analysis of telomeres (SMAT) assay (Sobinoff et al., 2017). The number of events and the length of the telomeric extension, which was marked by the incorporation of the thymidine analog 5-chloro-2’-deoxyuridine strictly at one end of the telomere, were increased in U2OS cells upon ATAD5 depletion (Fig. 4d-f). Taken together, these results suggest that ATAD5 depletion leads to telomere lengthening in ALT+ cancer cells.

**Fig. 4.**
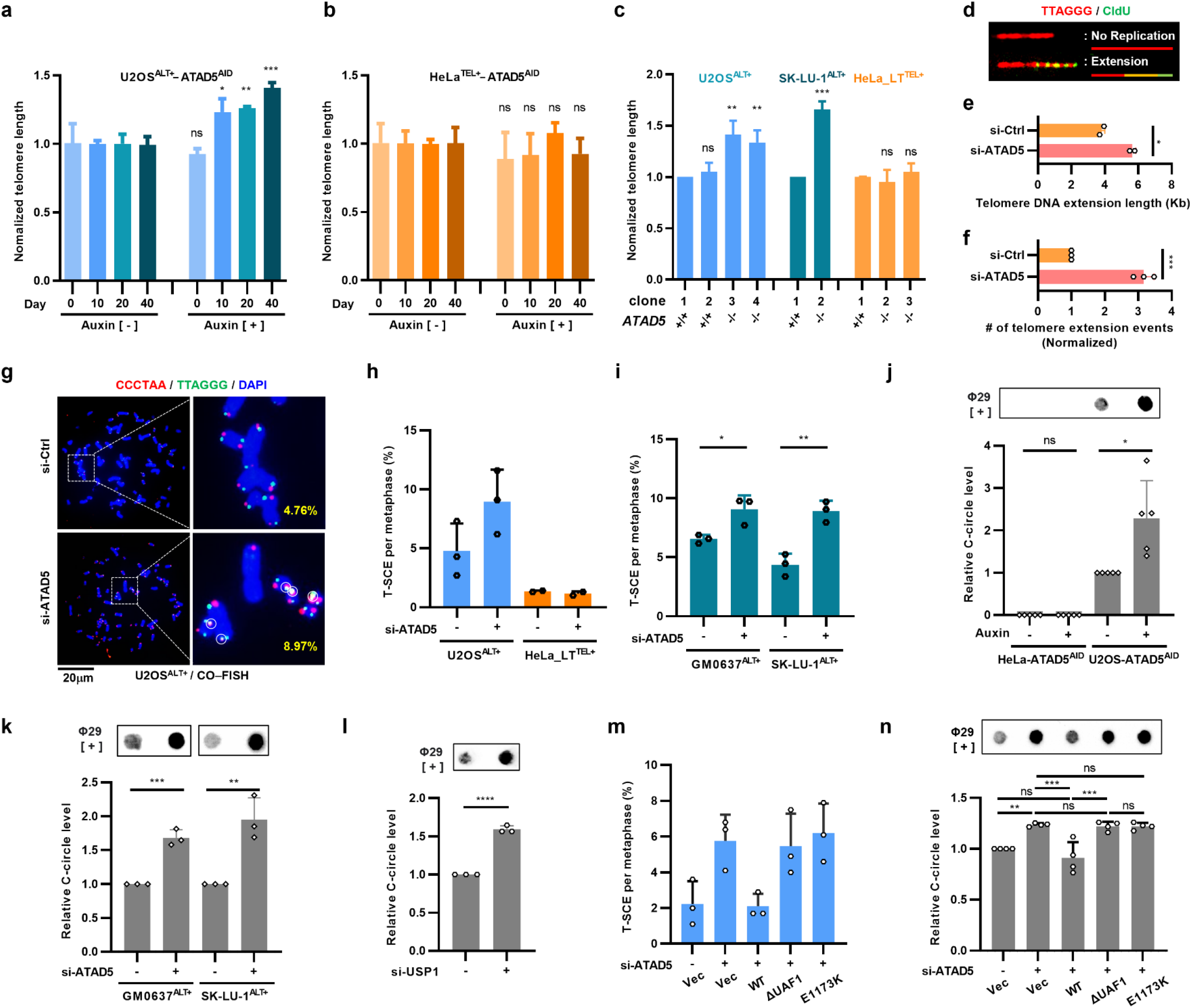
ATAD5 deficiency extends telomere length in ALT+ cancer cells and promotes recombination-associated ALT phenotypes. (a, b) U2OS-ATAD5^AID^ or HeLa-ATAD5^AID^ cells were treated with auxin for the indicated days and subjected to a Flow-FISH assay. (c) *ATAD5*^+/+^ and *ATAD5^−/−^* clones of each cell line were subjected to a Flow-FISH assay. (a-c) Normalized telomere length was quantified. (d-f) U2OS cells were transfected with siRNA and subjected to a SMAT assay. (d) Representative images of SMAT assay. (e, f) The extended telomere length (e) and the number of telomere extension events (f) were quantified. (g-I, m) After transfection, cells were subjected to a CO-FISH to measure T-SCE. (g) Representative images of CO-FISH. A circle indicates a T-SCE event. (h, i, m) The percentage of T-SCE per metaphase was quantified. (j) U2OS-ATAD5^AID^ or HeLa-ATAD5^AID^ cells were treated with auxin for 48 h. (j, k, l, n) The intensity of the C-circle level was quantified. (m, n) U2OS cells were transfected with a combination of *ATAD5* siRNA and wild type (WT), UAF1 interaction-defective (ΔUAF1) or PCNA unloading-defective (E1173K) ATAD5 cDNA. Data in all graphs represent the mean ± s.d. of at least three independent experiments. Statistical analysis: two-tailed unpaired Student’s *t*-test (a-c, e,f, i-l); One-way ANOVA (n).

### Ub-PCNA promotes recombination-associated ALT phenotypes

Telomeric DNA synthesis in APBs and MiDAS, both of which were increased upon ATAD5 depletion, involves a recombination process (Min et al., 2017; Zhang et al., 2019). Therefore, we investigated the effects of ATAD5 depletion on recombination-associated ALT phenotypes such as telomere-sister chromatid exchange (T-SCE) and C-circle, a type of extrachromosomal telomeric repeat that is a byproduct of telomere recombination in the ALT pathway (Cesare and Griffith, 2004). We found that ATAD5 depletion increased T-SCE and C-circle levels in three different ALT+ cell lines, but not in TEL+ HeLa-originated cells (Fig. 4g-k and Extended Data Fig. 4c-h). USP1 depletion also increased C-circle levels in U2OS cells (Fig. 4l). Consistently, the increases in T-SCE and C-circle induced by ATAD5 depletion were not restored by the UAF1-interaction-deficient ATAD5 (ATAD5^ΔUAF1^), which is defective in USP1-mediated PCNA deubiquitination (Lee et al., 2010), in contrast to the recovery to basal levels by wild-type ATAD5 (Fig. 4m, n and Extended Data Fig. 4i). Interestingly, the PCNA-unloading-defective ATAD5 ATPase mutant (ATAD5^E1173K^) showed similar defects (Fig. 4m, n). Taken together, these results suggest that both the PCNA deubiquitination and unloading activity of ATAD5 are required to inhibit recombination-associated ALT phenotypes.

### Unprocessed Ub-PCNA during the G2 phase increases APBs

Telomeric replication stress leads to APB formation and telomeric DNA synthesis in ALT+ cells (Cesare et al., 2009; Min et al., 2017). There is a possibility that ATAD5 or USP1 deficiency may have an effect on replication stress itself (Lee and Park, 2020). Therefore, we investigated the effect of ATAD5 or USP1 deficiency on ALT after excluding its influence on replication stress during the S phase. For this purpose, G2-phase synchronized U2OS-ATAD5^AID^ cells were treated with auxin for 6 h to acutely degrade the ATAD5 protein or treated with ML323, a chemical inhibitor of USP1-UAF1 (hereafter referred to as USP1i) (Liang et al., 2014). Both treatments increased chromatin PCNA, monoUb-PCNA, and diUb-PCNA levels (Fig. 5a and Extended Data Fig. 5a). The effects of USP1i on Ub-PCNA were more pronounced in the G2 phase (Extended Data Fig. 5a). Combining the two treatments further increased the levels of the monoUb- and diUb-PCNA (Fig. 5a). This could be explained by the persistence of all PCNA species on the chromatin due to the loss of the PCNA unloading activity of the ATAD5-RLC (Kang et al., 2019).

**Fig. 5.**
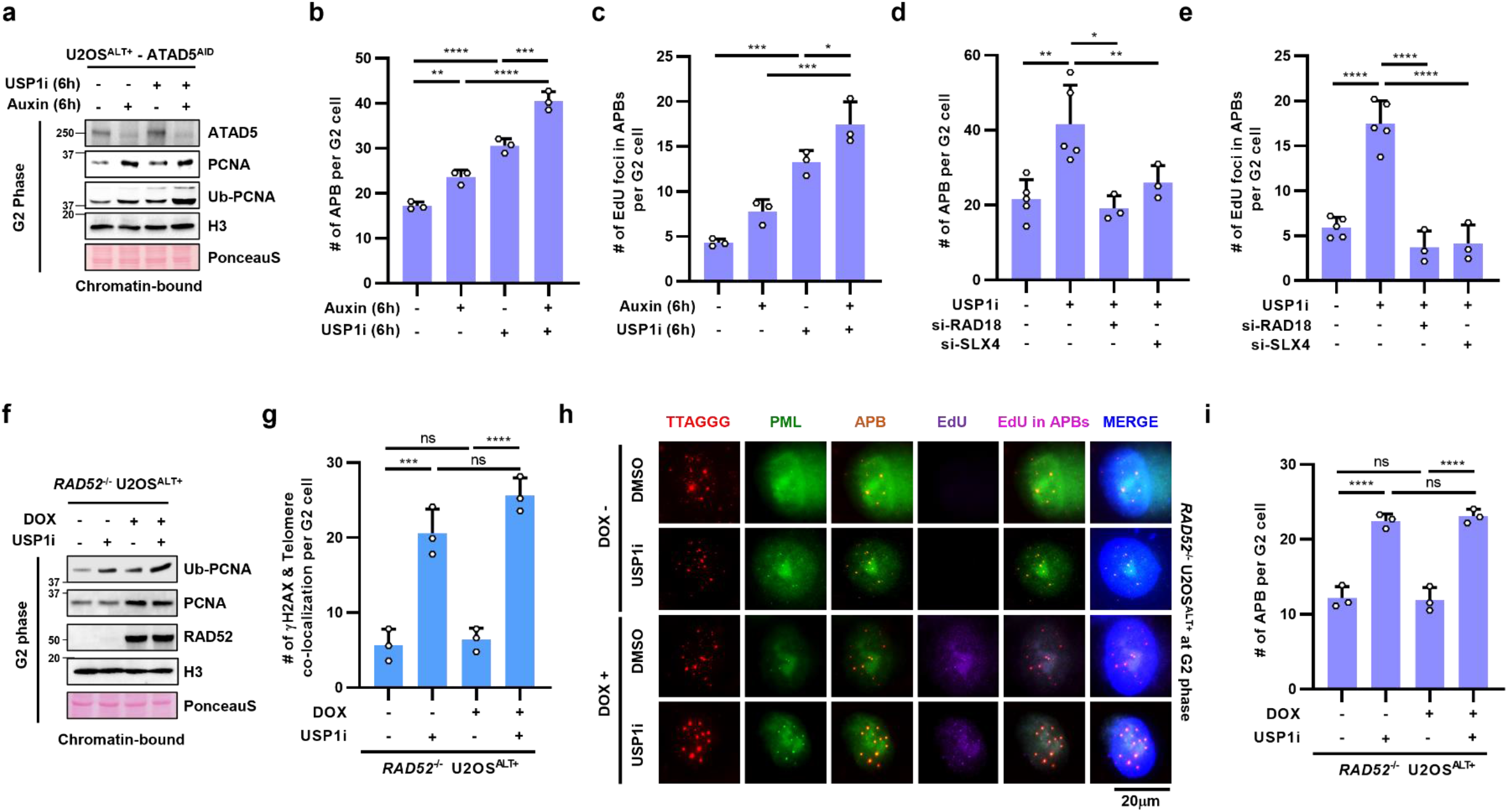
Ub-PCNA triggers telomere damage leading to APB formation and subsequent BIR in the G2 phase of ALT+ cancer cells. (a-c) U2OS-ATAD5^AID^ cells were synchronized in the G2 phase, treated with auxin and/or USP1i for 6 h, and chromatin-bound protein extracts were prepared for immunoblotting (a) or cells were subjected to an ATSA assay (b, c). (d, e) After transfection, U2OS cells were synchronized in the G2 phase, treated with USP1i for 6h, and subjected to an ATSA assay. (f-i) RAD52 doxycycline-inducible *RAD52*^−/−^ U2OS cells were synchronized in the G2 phase, treated with USP1i for 6 h, and chromatin-bound protein extracts were prepared for immunoblotting (f) or cells were fixed for γH2AX immunostaining to measure TIFs (g) or an ATSA assay (h, i). RAD52 expression was induced by doxycycline (DOX) treatment for 48 h before cell harvest or fixation. (h) Representative images of an ATSA assay. Data in all graphs represent the mean ± s.d. of at least three independent experiments. Statistical analysis: One-way ANOVA (b-e, g, i).

Acute ATAD5 degradation during the G2 phase increased the number of total APBs and EdU+ APBs (Fig. 5b, c). G2-specific USP1i treatment also increased the APBs and EdU+ APBs (Fig. 5b-e), which was further increased by ATAD5 depletion (Fig. 5b, c) but restored to basal levels when RAD18 was depleted (Fig. 5d, e and Extended Data Fig. 5b). This suggests that a substantial amount of Ub-PCNA is retained until the G2 phase to facilitate APB formation, which is promoted when Ub-PCNA is not deubiquitinated during the G2 phase.

### Unprocessed Ub-PCNA during the G2 phase induces telomere damage and APB formation in a RAD52-independent manner

Based on the increased number of APBs by USP1 inhibition, we speculated that the increase in Ub-PCNA levels may be involved in the generation of telomeric damage leading to APB formation. Therefore, we examined DNA damage foci at telomeres called telomere dysfunction-induced foci (TIFs) (Takai et al., 2003). The number of TIFs was increased in U2OS cells treated with USP1i in the G2 phase (Fig. 5f, g). The number of APBs in the G2 phase was minimally affected by RAD52 or POLD3 depletion (Zhang et al., 2019), suggesting that the clustering of damaged telomeres occurs before BIR initiation at APBs. Consistent with these results, there was no significant difference in TIFs induced by USP1i in wild type and *RAD52^−^*^/^*^−^* U2OS cells (Fig. 5f, g). As previously reported (Zhang et al., 2019), RAD52 deficiency completely inhibited DNA synthesis but did not affect total APB number (Fig. 5h, i). Interestingly, increased APB formation induced by USP1i was not affected by RAD52 deficiency (Fig. 5h, i). Taken together, these results suggest that unprocessed Ub-PCNA during the G2 phase can trigger telomere damage, leading to APB formation before subsequent BIR.

### The TLS polymerase REV1 is dispensable for increased APB numbers caused by unprocessed Ub-PCNA in the G2 phase

MonoUb-PCNA recruits Y-family TLS polymerases to bypass DNA lesions (Moldovan et al., 2007). REV1, a pivotal TLS polymerase, was reported to be required for MiDAS (Wu et al., 2023). We therefore investigated the involvement of TLS in the increased APB numbers caused by Ub-PCNA in the G2 phase. Treatment of U2OS cells in the G2 phase with a REV1 inhibitor (REV1i), JH-RE-06 (Wojtaszek et al., 2019), did not affect the number of total APBs and EdU+ APBs (Extended Data Fig. 5c, d). In addition, USP1i-induced APBs and EdU+ APBs were not affected by REV1i, suggesting that TLS is not involved in the increased APBs induced by unprocessed Ub-PCNA.

### SLX4 participates in Ub-PCNA-induced APB formation

SLX4 promotes MiDAS at CFSs and telomeres (Minocherhomji et al., 2015; Ozer et al., 2018). In addition, SLX4 is reported to be recruited to DNA damage sites via its ubiquitin-binding zinc finger 4 (UBZ4) domain (Katsuki et al., 2021; Kim et al., 2011; Lachaud et al., 2014). Because unprocessed Ub-PCNA due to ATAD5 depletion or USP1i treatment induces telomere damage and APB formation, we investigated whether SLX4 is involved in the process. Interestingly, we found that SLX4 depletion restored the USP1i-induced increases in APBs and EdU+ APBs to the levels similar to those achieved by RAD18 depletion in the G2 phase (Fig. 5d, e and Extended Data Fig. 5b).

### Ub-PCNA increases SLX4 recruitment to telomeres in ALT+ cells, but not in TEL+ cells in the G2 phase

Since SLX4 is involved in ALT regulation, we next investigated the effects of Ub-PCNA on telomeric localization of SLX4 in U2OS cells. As previously reported (Wan et al., 2013; Wilson et al., 2013), a portion of SLX4 foci colocalized with telomeres through the TRF2 binding motif (TBM) in the G2 phase (Fig. 6a, b). The localization of SLX4 at telomeres was increased by ATAD5 depletion but decreased by RAD18 depletion.

**Fig. 6.**
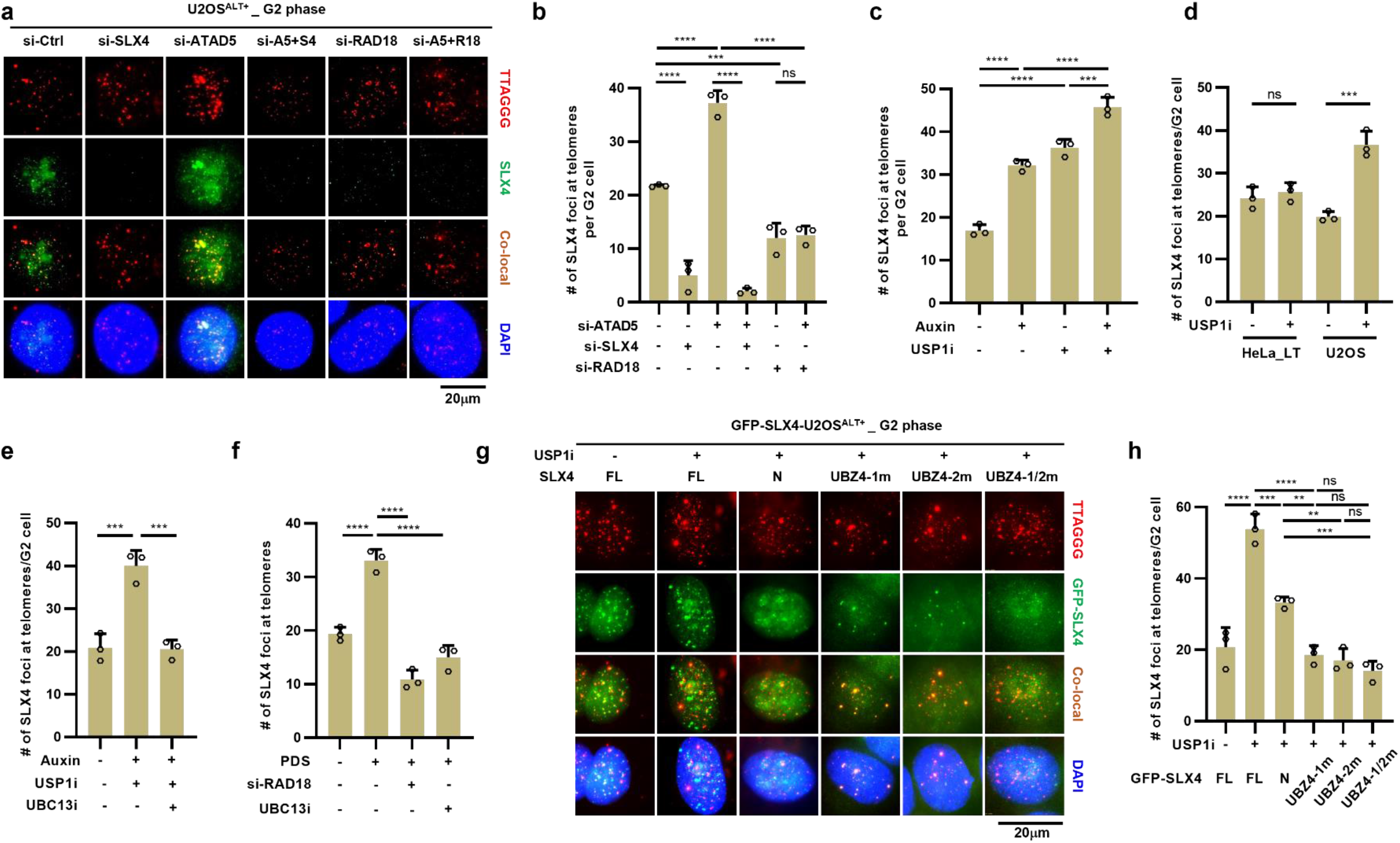
PolyUb-PCNA increases SLX4 recruitment to telomeres in ALT+ cancer cells. (a, b) After transfection, U2OS cells were synchronized in G2 phase and fixed for SLX4 immunostaining with telomere FISH. (a) Representative images showing SLX4 localization at telomeres. (b-f, h) The number of SLX4 localization at telomeres in the G2 phase was quantified. (c) U2OS-ATAD5^AID^ synchronized in the G2 phase were treated with auxin and USP1i for 6 h and fixed. (d) HeLa_LT or U2OS cells synchronized in the G2 phase were treated with USP1i for 6 h and fixed. (e) G2-synchronized U2OS-ATAD5^AID^ cells were treated with auxin, USP1i, and UBC13i for 6 h, and cells were fixed. (f) After siRNA transfection, U2OS cells were treated with pyridostatin (PDS) for 24 h and UBC13i for 6 h, respectively, before fixation. (g, h) U2OS cells expressing GFP-tagged full-length SLX4 (FL), N-terminal fragment of SLX4 (N), SLX4^N^ with a mutation in UBZ4-1 (UBZ4-1m), UBZ4-2 (UBZ4-2m), or both UBZ4 domains (UBZ4-1/2m), respectively, were synchronized in the G2 phase, treated with USP1i for 6 h and fixed. (g) Representative images showing GFP-SLX4 localization at telomeres. Data in all graphs represent the mean ± s.d. of at least three independent experiments. Statistical analysis: One-way ANOVA (b, c, e, f, h); two-tailed unpaired Student’s *t*-test (d).

Furthermore, the enhanced localization of SLX4 at telomeres induced by ATAD5 depletion was reduced by RAD18 co-depletion. G2-specific ATAD5 degradation or USP1i treatment also increased SLX4 localization at telomeres in U2OS-ATAD5^AID^ cells (Fig. 6c). In contrast, telomeric SLX4 localization was not affected by G2-specific USP1 inhibition in HeLa_LT cells (Fig. 6d and Extended Data Fig. 6a). Taken together, these results suggest that unprocessed Ub-PCNA during the G2 phase facilitates SLX4 recruitment to telomeres in ALT+ cells.

### SLX4 is recruited to telomeres containing polyUb-PCNA in a UBZ4 domain-dependent manner

SLX4 is recruited to interstrand crosslink lesions by interacting with K63-linked polyubiquitin chains through its UBZ4 domain (Katsuki et al., 2021; Kim et al., 2011; Lachaud et al., 2014). Since UBC13-mediated PCNA polyubiquitination is also K63-linked (Lee and Myung, 2008), we determined whether SLX4 recruitment to telomeres requires polyUb-PCNA by observing SLX4 recruitment to telomeres upon downregulating PCNA deubiquitination. We first combined G2-specific ATAD5 degradation with USP1i treatment using U2OS-ATAD5^AID^ cells to increase chromatin polyUb-PCNA levels. Under these conditions, treatment with a UBC13 inhibitor (UBC13i), NSC697923 downregulated PCNA polyubiquitination (Pulvino et al., 2012) (Extended Data Fig. 6b) and reduced SLX4 localization at telomeres to basal levels (Fig. 6e), strongly suggesting that polyUb-PCNA recruits SLX4 to telomeres in ALT+ cells.

Telomeric G4 signals are frequently found in ALT+ cancer cells and G4 stabilization promotes ALT activity (Amato et al., 2020; Min et al., 2017; Yang et al., 2021a). We found that a G4 stabilizer pyridostatin (PDS) treatment increased both monoUb- and diUb-PCNA levels on chromatin in two different ALT+ cancer cell lines (Extended Data Fig. 6c). The colocalization of SLX4 foci with telomeres was also increased by PDS treatment, which was reduced to basal levels by RAD18 depletion or UBC13i treatment (Fig. 6f and Extended Data Fig. 6d), suggesting sustained G4s recruit SLX4 to telomeres via PCNA polyubiquitination in ALT+ cancer cells.

Next, we investigated the role of the UBZ4 domain of SLX4 in its recruitment to telomeres with unprocessed Ub-PCNA during the G2 phase. We used U2OS cells expressing the following SLX4 constructs, each fused to a green fluorescent protein (GFP) (Katsuki et al., 2021): full-length SLX4 (SLX4^FL^), N-terminal fragment of SLX4 (SLX4^N^), and SLX4^N^ with a mutation in the UBZ4-1 (SLX4^UBZ4-1m^), UBZ4-2 (SLX4^UBZ4-2m^), or both UBZ4 domains (SLX4^UBZ4-1/2m^) (Fig. 6g, h and Extended Data Fig. 7a). Protein expression levels and foci numbers in the G2 phase were comparable among SLX4^N^ and the three SLX4^N^ mutants (Extended Data Fig. 7b, c). USP1 inhibition increased SLX4^FL^ localization at telomeres in U2OS cells (Fig. 6g, h). SLX4^N^ exhibited less localization at telomeres than SLX4^FL^, probably due to the lack of a TBM domain. Interestingly, SLX4^UBZ4-1m^, SLX4^UBZ4-2m^, and SLX4^UBZ4-1/2m^ showed significantly reduced localization at telomeres compared with SLX4^N^ under USP1i-treated conditions (Fig. 6g, h). Taken together, these data suggest that SLX4 is recruited to polyUb-PCNA at telomeres through its UBZ4 domain.

**Fig. 7.**
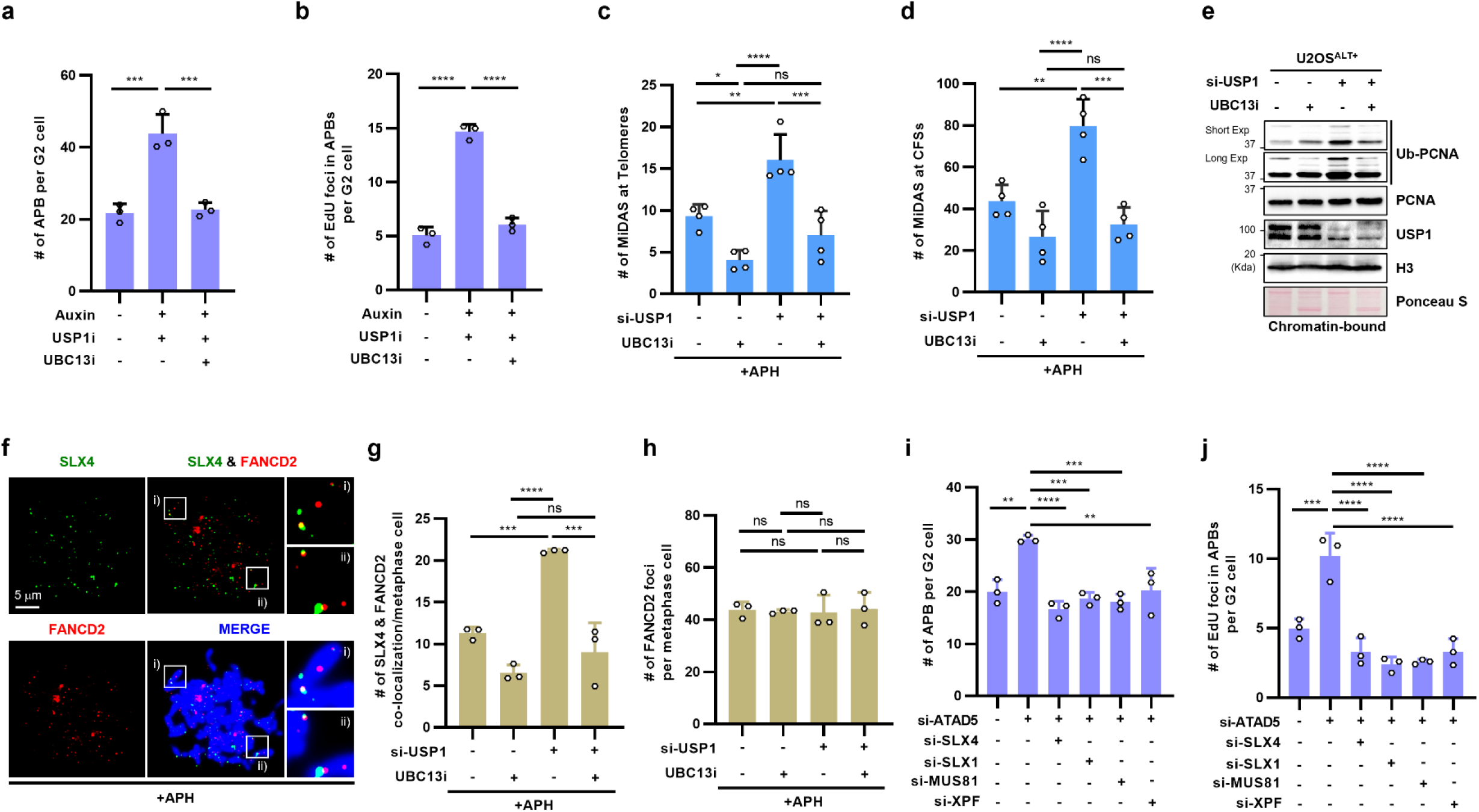
PolyUb-PCNA induces APB formation and MiDAS at telomeres and CFS. (a, b) G2-synchronized (G2) U2OS-ATAD5^AID^ cells were treated with auxin, USP1i, and UBC13i for 6 h, and cells were fixed for an ATSA assay. (c-h) After transfection, U2OS cells were subjected to a MiDAS assay (c, d), immunoblotting with chromatin-bound protein extracts (e), or immunostaining (f-h). (c, d, f-h) U2OS cells were treated with 0.4 μM aphidicolin (APH) for 16 h, and 10 μM UBC13i for 17 h before fixation as indicated. (c, d) The number of MiDAS at telomeres (c) or CFSs (d) was quantified. (f) Representative images showing SLX4 and FANCD2 localization in metaphase cells. (g, h) The number of SLX4 and FANCD2 co-localization (g) and FANCD2 foci (h) per metaphase cell was quantified. (i, j) After transfection, U2OS cells synchronized in the G2 phase were fixed for an ATSA assay. Data in all graphs represent the mean ± s.d. of at least three independent experiments. Statistical analysis: One-way ANOVA (a-d, g-j).

### Unprocessed polyUb-PCNA induces APB formation and MiDAS at telomeres and CFS

Consistent with the role of polyUb-PCNA in SLX4 recruitment to telomeres, UBC13 inhibition restored the increases in APBs including EdU+ APBs induced by G2-specific ATAD5 degradation and USP1i treatment in U2OS-ATAD5^AID^ cells (Fig. 7a, b). These results strongly suggest that the recruitment of SLX4 to telomeres by polyUb-PCNA is responsible for telomere damage and subsequent APB formation in ALT+ cells.

MiDAS at telomeres and CFS requires SLX4 (Minocherhomji et al., 2015; Ozer et al., 2018). Inspired by the findings that polyUb-PCNA is involved in the recruitment of SLX4 for BIR initiation at telomeres, we investigated the involvement of polyUb-PCNA in MiDAS. UBC13 inhibition alone reduced MiDAS at both telomeres and CFSs in U2OS cells (Fig. 7c-e). Consistently, UBC13 inhibition reversed the increased MiDAS by USP1 depletion (Fig. 7c-e). Furthermore, UBC13 inhibition reduced SLX4 localization at FANCD2 foci, which are associated with CFS expression in metaphase (Chan et al., 2009), and restored the increases induced by USP1 depletion in U2OS cells (Fig. 7f, g), while the number of FANCD2 foci remained constant (Fig. 7h). Taken together, these results suggest that polyUb-PCNA-mediated SLX4 recruitment is responsible for MiDAS at telomeres and CFS in ALT+ cells.

### SLX4-interacting nucleases facilitate APB formation and telomeric DNA synthesis in ATAD5-deficient ALT+ cells

SLX4 is a scaffold for several nucleases such as SLX1, MUS81-EME1, and XPF-ERCC1 (Munoz et al., 2009). We investigated which nuclease(s) are responsible for the increased ALT activity associated with unprocessed Ub-PCNA in ALT+ cells. Unexpectedly, depletion of any of the three SLX4-interacting nucleases restored the increases in total APBs and EdU+ APBs induced by ATAD5 depletion to levels similar to those caused by SLX4 co-depletion (Fig. 7i, j and Extended Data Fig. 7d, e). These results suggest that SLX1, MUS81-EME1, and XPF-ERCC1 cooperatively generate DNA breaks at telomeres associated with polyUb-PCNA, leading to APB formation and ALT processes.

## Discussion

PCNA ubiquitination plays a crucial role in directing damage bypass pathway and occurs during DNA replication and DNA repair synthesis (Hoege et al., 2002; Motegi et al., 2008; Unk et al., 2008; Zlatanou et al., 2011). Our study reveals the role of Ub-PCNA in BIR-associated ALT processes during the G2 phase. We propose that polyUb-PCNA contributes to these processes by inducing telomere breakage and facilitating RAD52-dependent BIR [Extended Data Fig. 8a (i)-(iii)]. We found that polyUb-PCNA at telomeres recruits SLX4 and associated nucleases, leading to telomere breaks and APB formation. Inhibition of PCNA polyubiquitination reduced APB levels and telomeric MiDAS events in ALT+ cells. Suppression of USP1-mediated PCNA deubiquitination during the G2 phase increased TIF signals and APB levels specifically in ALT+ cells in a RAD18-, UBC13-, and SLX4-dependent manner. Consistently, ATAD5 or USP1 deficiency increased telomeric MiDAS events in a RAD18-, UBC13-, and RAD52-dependent manner. Unprocessed Ub-PCNA during the G2 phase increased telomeric SLX4 localization, dependent on RAD18, UBC13 and SLX4 UBZ4 domain. Excessive telomeric DNA breakage and recombination caused by uncontrolled polyUb-PCNA-SLX4 activity can lead to telomere instability [Extended Data Fig. 8b (i)]. Therefore, this activity must be limited to tolerable levels, and this counterbalance likely relies on proper deubiquitination and/or unloading of polyUb-PCNA by the USP1-UAF1-ATAD5-RLC [Extended Data Fig. 8a (ii)]. Additionally, we propose that Ub-PCNA enhances the BIR process in ALT+ cells [Extended Data Fig. 8a (iv)], as observed through its localization at telomeric break sites and the promotion of break-induced telomere DNA synthesis. Overall, Ub-PCNA serves as both an inducer of telomere breakage and a promoter of the BIR process in BIR-associated ALT processes.

We observed RAD18- and UBC13-dependent MiDAS events not only at telomeres but also at CFSs in ALT+ U2OS cells. UBC13 inhibition decreased SLX4 localization at FANCD2 foci (sites of CFS expression) in metaphase, while USP1 depletion increased it in a UBC13-dependent manner, similar to telomeres. We speculate that the molecular mechanisms associated with polyUb-PCNA are primarily utilized to alleviate replication stress of telomeres, potentially leading to an increase in replication stress at CFSs in ALT+ cells. In addition, CFS, which has longer replication times and is sensitive to replication stress (Helmrich et al., 2011), may be particularly dependent on the polyUb-PCNA-associated BIR mechanism. Interestingly, a high-resolution mapping study in U2OS cells showed that there are additional MiDAS regions that do not overlap with HeLa cells (Macheret et al., 2020), possibly corresponding to CFSs regulated by polyUb-PCNA-associated BIR mechanisms in U2OS cells.

Our study found that ALT+ cancer cells exhibited increased MiDAS events and recombination-associated ALT phenotypes due to unprocessed Ub-PCNA, while TEL+ HeLa_LT cells did not. The ALT cell-specific effect may be attributed to the higher levels of replication stress inherent in ALT telomeres (Lu and Pickett, 2022). We observed higher monoUb-PCNA levels in ALT+ U2OS cells compared to TEL+ HeLa_LT cells, and polyUb-PCNA was only present in U2OS cells, particularly after G2-specific USP1 inhibition (Extended Data Fig. 5a, 6a). These differences in polyUb-PCNA levels may lead to SLX4-mediated telomeric breakage specifically in ALT+ cells. The presence of APBs, which contain various recombination and replication proteins (Draskovic et al., 2009; Min et al., 2017; Zhang et al., 2019), in ALT+ cells may provide opportunities for RAD52-mediated BIR that can be triggered by ATAD5 deficiency or USP1 inhibition. In addition, ALT telomeres, which are characterized by greater replication difficulties, may require more Ub-PCNA for the BIR process compared to TEL telomeres. For these reasons, we suggest that the effects of Ub-PCNA in promoting telomeric BIR and ALT phenotypes may be specific to ALT+ cells.

The PCNA-unloading-defective ATAD5^E1173K^ exhibited increased break- and recombination-associated ALT phenotypes similar to ATAD5 depletion (Fig. 4), which indicates that the PCNA unloading activity of ATAD5 is necessary to limit ALT phenotypes. ATAD5^E1173K^ can facilitate PCNA deubiquitination, but it is unclear whether the PCNA promoting ALT phenotypes in cells expressing ATAD5^E1173K^ is unmodified or ubiquitinated. Ub-PCNA remains on chromatin even in the G2 phase (Extended Data Fig. 5a), and monoUb-PCNA-dependent TLS levels are higher in the G2 phase compared to the S phase (Diamant et al., 2012). This suggests that there is a mechanism to restrict the deubiquitination and unloading of Ub-PCNA before completing its task in the G2 phase. Therefore, we speculate that Ub-PCNA, rather than unmodified PCNA, present at telomeres in the G2 phase contributes to the augmented ALT phenotypes in cells expressing ATAD5^E1173K^.

SLX4 localizes at telomeres by interacting with TRF2 through its TBM domain, preventing telomere damage and negatively regulating telomere length in TEL+ and ALT+ cancer cells (Wan et al., 2013; Wilson et al., 2013). SLX4 also counteracts BLM- and POLD3-mediated telomere synthesis and ALT activity through its TBM domain in ALT+ cancer cells (Sobinoff et al., 2017). However, in the present study, a positive role of SLX4 in BIR-associated ALT processes was observed, which depended on its UBZ4 domain rather than the TBM domain. The recruitment of SLX4 to interstrand crosslink lesions occurs through its UBZ4 domain, which interacts with K63-linked polyubiquitin chains on proteins around the lesions (Katsuki et al., 2021; Kim et al., 2011; Lachaud et al., 2014). We propose that K63-linked polyUb-PCNA at telomeres serves as a binding platform for SLX4 and its associated SMX tri-nuclease complex (SLX1, MUS81-EME1, and XPF-ERCC1), leading to telomere breakage in the G2 phase. Although each structure-specific endonucleases (SLX1, MUS81, and XPF) in the SMX complex has different substrate preferences (Wyatt et al., 2017), all three nucleases are required for telomere breakage caused by polyUb-PCNA-SLX4 (Fig. 7i, j). This requirement for all three nucleases was also observed in T-SCE in U2OS cells (Wan et al., 2013). The specific telomeric DNA structures associated with polyUb-PCNA are currently unclear, but it appears that they must be processed sequentially or cooperatively by the entire SMX tri-nuclease complex.

## Acknowledgments

We thank Dr. Roger A. Greenberg, Dr. Lee Zou, Dr. Jan Karlseder, and Dr. Minoru Takata for generously providing and sharing technical information about the U2OS-mCherry-TRF1-FokI cells, the RAD52 inducible U2OS-*RAD52*^−/−^ (clone #2) cells, the HeLa_LT cells, and the CSII-based SLX4 DNA constructs, respectively. We also thank RIKEN BRC for the MTA agreement (#DNA86581) on the CSII-based SLX4 DNA constructs. We sincerely thank members of the Center for Genomic Integrity, IBS for helpful discussions and comments. S.K. is thankful for the support of the ASAN Foundation Biomedical Science scholarship.

## Author contributions

K.L. and K.M. conceptualized the project. S.K. and K.L. designed the experiments. S.K. performed and analyzed most cell-based experiments. N.K. performed and analyzed most T-SCE experiments. J.S.R. performed and analyzed the SMAT assay and part of the T-SCE experiments. S.P. generated part of the cell line. K.L. and K.M. supervised the project. S.K. and K.L. wrote the manuscript with help from all other authors.

## Funding

This research was supported by the Institute for Basic Science (IBS-R022-D1) and National Research Foundation of Korea (NRF) grant funded by the Korea government (MIST) (RS-2023-00251939 to K.L).

## Methods

### Cell lines and cell culture

HeLa, HeLa-LT (kindly provided by Dr. Jan Karlseder), GM0637, SK-LU-1, U2OS, U2OS-ATAD5^AID^ (Park et al., 2019) and HeLa-ATAD5^AID^ (Park et al., 2021) cells were cultured in DMEM (Hyclone) supplemented with 10% fetal bovine serum (Hyclone), 100 U/mL penicillin G (Life Technologies), and 100 mg/mL streptomycin (Life Technologies) in a humidified atmosphere of 5% CO_2_ at 37°C. The RAD52-inducible *RAD52*^−/−^ U2OS cells (Zhang et al., 2019) (kindly provided by Dr. Lee Zou) and U2OS-Tet-On-destabilization domain (DD)-estrogen receptor (ER)-mCherry-TRF1-FokI cells (U2OS-mCherry-TRF1-FokI) cells (Dilley et al., 2016) (kindly provided by Dr. Roger A. Greenberg) were cultured in DMEM supplemented with 10% tetracycline-free fetal bovine serum (Takara) and antibiotics. RAD52 protein expression was induced in *RAD52*^−/−^ U2OS cells by treating with 100 ng/mL doxycycline for 48 h. The *ATAD5*^−/−^ U2OS, *ATAD5*^−/−^ SK-LU-1, and *ATAD5*^−/−^ HeLa-LT cell lines were generated using commercial ATAD5 CRISPR/Cas9 knockout plasmids (sc-405654) (Santa Cruz). Briefly, cells were transfected with the CRISPR/Cas9 plasmids, and, after 48 h transfection, GFP-positive cells were sorted using a FACSAria cell sorter (BD biosciences). After 3 weeks, single cell colonies were picked, and complete *ATAD5* knockout was confirmed by immunoblotting. The U2OS cells expressing GFP-tagged full-length SLX4, N-terminal fragment of SLX4 (SLX4^N^), and SLX4^N^ with a mutation in UBZ4-1, UBZ4-2, or both UBZ4 domains, respectively, were generated using the CSII-CMV-based lentiviral vectors reported in a previous study (Katsuki et al., 2021). Briefly, the lentiviral vectors encoding each SLX4 cDNA (kindly provided by Dr. Minoru Takata) were individually co-transfected with two assistant vectors pMD2.G (#12259, Addgene) and psPAX2 (#12260, Addgene) into HEK293-FT cells. After 48 h, viral supernatants were collected and clarified by using a 0.2 μm filter. The lentivirus was then infected into U2OS cells with polybrene (sc-134220, Santacruz) followed by selection with 3 μg/ml of blasticidin (R21001, Thermo Fisher) for 5 days. Transfection of plasmid DNA was performed using X-tremeGENE™ HP (Roche) and siRNAs were transfected using RNAiMAX (Thermo Fisher). Cells were analyzed 48 h after transfection.

### TRF1-FokI-induced generation of telomeric breaks in the G2 phase

To generate telomeric breaks in U2OS-mCherry-TRF1-FokI cells in the G2 phase, cells were simultaneously treated with 100 ng/mL doxycycline (induces expression of DD-ER-mCherry-TRF1-FokI proteins) and 15 μM RO-3306 (arrests cells in the G2 phase) for 16 h, and then treated with 1 μM tamoxifen (binds to ER and induces nuclear transport of proteins fused to ER) and 1 μM Shield1 (stabilizes proteins tagged with a DD domain) for 2 h, otherwise descripted in the figure legends. To detect break-induced telomere synthesis (EdU localization at TRF1-FokI), 10 μM EdU was added to cells before 10 min fixation.

### siRNAs

The following synthetic duplex siRNAs were purchased from Bioneer: ATAD5 3ʹ UTR (5ʹ-GUAUA UUUCU CGAUG UACA-3ʹ), RAD52 (5ʹ-AGACU ACCUG AGAUC ACUAtt-3ʹ), POLD3 (5ʹ-GGCCU CUGUU CAAUA CUGAtt-3ʹ), RAD18 (5ʹ-GCUCU CUGAU CGUGA UUUA-3ʹ), USP1 (5ʹ-AGCUA CAAGU GAUAC AUUAtt-3ʹ), SLX4 5ʹ UTR [SLX4 UTR87 (5ʹ-GCACC AGGUU CAUAU GUAUtt-3ʹ)], SLX4 3ʹ UTR [SLX4 UTR7062 (5ʹ-GCACA AGGGC CCAGA ACAAtt-3ʹ)], SLX1 (5ʹ-UGGAC AGACC UGCUG GAGAU Utt-3ʹ), MUS81 (5ʹ-UAGGA UCUUA GCUUC CUUCC CGCUGtt-3ʹ), XPF (5ʹ-GUAGG AUACU UGUGG UUGAtt-3ʹ), and control siRNA (Bioneer #SN-1002). RNF168 siRNA (5ʹ-GACAC UUUCU CCACA GAUAU U-3ʹ) was purchased from GENOLUTION. SLX4 5ʹ UTR- and 3ʹ UTR-targeting siRNAs were used as a mixture.

### Chemicals, reagents and antibodies

The following chemicals were used in this study: indole-3-acetic acid (IAA, auxin) (87-51-4) (Millipore); 4-hydroxytamoxifen (4-OHT, tamoxifen) (H7904), aphidicolin (A0781), thymidine (T1895), RO-3306 (SML0569, a CDK1 inhibitor), 5-ethynyl-2ʹ-deoxyuridine (EdU) (#900584), 5-bromo-2ʹ-deoxyuridine (BrdU) (B5002), doxycycline hyclate (doxycycline) (D9891) (Sigma-Aldrich); 5-bromo-2ʹ-deoxycytidine hydrate (BrdC) (#0210016680), 5-chloro-2ʹ-deoxyuridine (CldU) (#105478) (MP Biomedicals); Shield1 (632189) (TAKARA); colcemid (15212-012) (Gibco); ML323 (S7529, an USP1-UAF1 inhibitor, 30 μM), JH-RE-06 (S8850, a REV1 inhibitor, 2.5 μM), NSC697923 (S7142, UBC13-UEV1A inhibitor, 10 μM), pyridostatin (PDS) (S7444, G4 stabilizer, 10 μM) (Selleckchem). The following reagents were used in this study: Hoechst 33258 (H1398), Ultrapure 20X SSC (#15557044) (Invitrogen); DIG Easy Hyb solution (#11603558001), Ponceau S solution (P7170) (Sigma-Aldrich); Exonuclease III (M1815), Set of each dNTPs (U1130) (Promega); formaldehyde (P2031), formamide (FC1014) (Biosesang). The following antibodies were used: anti-UAF1 (sc-514473), anti-GFP (sc-8334), anti-RAD52 (sc-365341), anti-PML (sc-966) (Santa Cruz Biotechnology); rabbit-anti-PCNA (ab18197), anti-RAD18 (ab57447), horseradish peroxidase (HRP)-conjugated anti-Digoxigenin (HRP.21H8) (ab6212), anti-FANCD2 (ab2187) for immunoblotting, anti-FANCD2 (ab108928) for immunostaining, anti-SLX4 (ab169114) for immunoblotting, anti-MUS81 (ab14387), anti-CldU (ab6326) (Abcam); anti-FLAG (F7425) (Sigma-Aldrich); anti-ubiquityl-PCNA (Lys164) (13439S), anti-Cyclin B1 (4138S), anti-histone H3 (#9175) (Cell signaling); anti-γH2AX (05-636) (Merck Millipore); anti-POLD3 (H00010714-M01) (Abnova); anti-USP1 (A301-698A), anti-PML (A301-167A) for immunostaining (Bethyl); anti-ATRX (GTX101310) (GeneTex); anti-XPF (NBP2-58407) (Novus Biologicals); anti-SLX4 (DU16029) (MRC laboratory for immunostaining) antibodies. The anti-human ATAD5 antibody was raised in rabbits against an N-terminal fragment (1–297 aa) (Lee et al., 2010).

### Reverse transcription-quantitative real-time PCR (RT-qPCR)

Total RNA was extracted using a TRIzol® Reagent (Invitrogen) protocol. 1.5 μg RNA was used to synthesize cDNA using the SuperScript® IV First-Strand Synthesis System (18091050) (Invitrogen). RT-qPCR was performed using Power SYBR Green master mix (4368702) (Applied Biosystems) in a QuantStudio 7 Flex system (Applied Biosystems) according to the manufacturer’s instructions. Gene expression was normalized to *36B4* expression. The following primers were used: 36B4U-forward (For) (5ʹ-CAGCA AGTGG GAAGG TGTAA TCC-3ʹ), 36B4D-reverse (Rev) (5ʹ-CCCAT TCTAT CATCA ACGGG TACAA-3ʹ), RNF168-For (5ʹ-GGCGA GTTTA TGCTG TCCCT-3ʹ), RNF168-Rev (5ʹ-GCCGC CACCT TGCTT ATTTC-3ʹ), SLX1-For (5ʹ-ATGCC CTTGC TGTGA GAAGT-3ʹ), SLX1-Rev (5ʹ-GGTAG GAGGG GGTAA GGACA-3ʹ).

### Protein extraction and immunoblot analysis

Protein extraction and immunoblot analysis were performed as previously described (Kim et al., 2020). For whole-cell proteins extraction, cell pellets were lysed in RIPA buffer (50 mM Tris-HCl, pH 8.0, 150 mM NaCl, 5 mM EDTA, 1% Triton X-100, 0.1% sodium dodecyl sulfate, 0.5% sodium deoxycholate, 0.1 M phenylmethylsulfonyl fluoride, phosphatase inhibitors, and protease inhibitors) with Benzonase nuclease for 45 min at 4°C, followed by sonication and centrifugation. For chromatin-bound protein extraction, cell pellets were first resuspended in Buffer A (100 mM NaCl, 300 mM sucrose, 3 mM MgCl_2_, 10 mM PIPES, pH 6.8, 1 mM EGTA, 0.2% Triton X-100, 0.1 M phenylmethylsulfonyl fluoride, phosphatase inhibitors, and protease inhibitors) and incubated for 5 min at 4°C. After centrifugation, pellets were further lysed in RIPA buffer. For immunoblot analysis, proteins were separated by SDS-PAGE and transferred to a nitrocellulose membrane. The blot was blocked with Tris-buffered saline containing 0.1% Tween 20 (TBST) supplemented with 5% skim milk for 30 min and incubated with primary antibodies overnight. After washing with TBST buffer, the blot was incubated with HRP-conjugated secondary antibodies (Enzo Life Sciences) for 30 min. After washing with TBST buffer, the signal was detected using an enhanced chemiluminescence reagent (Thermo Fisher Scientific) with an automated imaging system (ChemiDoc™; Bio-Rad Laboratories).

### Immunostaining

Immunostaining with the metaphase spread was performed as previously described (Kim et al., 2020) with slight modification. After trypsinization, cells were incubated with hypotonic solution (75 mM KCl) and fixed with fixative solution (methanol/acetic acid 3:1). Fixed cells were spread onto a slide and incubated with pre-extraction buffer (0.1% Triton X-100, 20 mM HEPES-KOH, pH 7.9, 50 mM NaCl, 3 mM MgCl_2_, 300 mM sucrose) for 10 min at 4°C. The slide was then rinsed with 1X cold phosphate-buffered saline (PBS), and cells were fixed with 4% paraformaldehyde (PFA) for 15 min and permeabilized with 0.5% Triton X-100 in PBS for 5 min. Cells were then blocked with blocking buffer [5% bovine serum albumin (BSA) and 0.5% Triton X-100 in PBS] for 1 h, and incubated with the primary antibody in blocking solution overnight at 4°C. After washing with the wash buffer (0.1% Triton X-100 in PBS), cells were incubated with Alexa Fluor-conjugated secondary antibody (1:500) in wash buffer for 1 h at room temperature. After washing with washing buffer, the cells were mounted using ProLong® Gold antifade reagent with DAPI (H-1200) (Vector Laboratories) or proceeded for further assays.

### Telomere fluorescence *in situ* hybridization (Telomere FISH)

Telomere FISH was performed after immunostaining and/or EdU-Click reaction. The cells in the slide were blocked by blocking solution (3% BSA in PBST). Cells were then fixed with 4% PFA for 10 min, dehydrated by sequential incubation in 75%, 85%, and 100% ethanol for 2 min each, and allowed to air dry completely. For probe denaturation, hybridizing solution (70% formamide, 12 mM Tris-pH 7.5, 5 mM KCl, 1 mM MgCl_2_ in 2X SSC) with 8 μg/ml peptide nucleic acid (PNA) probes TelC (repeats of CCCTAA)-Cy3 (F1002, Panagene) or TelG (repeats of TTAGGG)-FITC (F1010, Panagene) was preheated to 72°C for 5 min and immediately cooled at 4°C for 5 min. Air-dried cells were incubated in warmed 70% formamide solution in 2X SSC at 68°C for 5 min to denature genomic DNA. Cells were then incubated with hybridizing solution containing denatured telomere probes at 37°C for 2 h in the dark. Cells were washed at room temperature with 50% formamide solution for 15 min, 2X SSC buffer, and 0.1% NP-40 in 2X SSC solution.

### Single-molecule analysis of telomeric DNA (SMAT) assay

SMAT assay was performed as previously described (Margalef et al., 2018) with slight modification. Briefly, cells were labeled with 100 μM CIdU for 4 h before harvest. Cells were embedded in 1.2% low-melting agarose (#1613112, Bio-Rad) plugs and then subjected to digestion with MboI restriction enzyme (R0147M, NEB) and proteinase K (#3115828001, Roche) overnight. Plugs were then dissolved with β-Agarase I (M0392S, NEB). Molecular combing was performed using the Molecular Combing System (Genomic Vision) with a constant stretch factor of 2 kb/μm using vinyl silane coverslips. Coverslips were then dried for 4 h at 65°C, and DNA was denatured in 2.5 M HCl for 1 h. DNA was then subjected to immunostaining for CldU and followed telomere FISH with TelG-Cy3 probe (F1006, Panagene).

### Chromosome orientation-FISH (CO-FISH)

CO-FISH assay to detect T-SCE was performed as previously described (Williams and Bailey, 2009) with slight modification. Cells were incubated with 10 μM of BrdU:BrdC (3:1) for 24 h, for 18 h in the case of the GM0637 cell line, before fixation. For removal of BrdU- and BrdC-incorporated (newly synthesized) DNA strand, the slide was incubated in RNase A (100 μg/ml) at 37°C for 10 min. DNA was then stained with Hoechst 33258 (500 μg/ml) in 2X SSC at room temperature for 15 min followed by a rinse with triple distilled water (TDW) and air dry. The slide was exposed to 365 nm ultraviolet light for 30 min and incubated with Exonuclease III solution (10 U/μl) at room temperature for 30 min. Telomere FISH was performed twice in a row by using TelG-FITC and TelC-Cy3 probes.

### ALT telomere DNA synthesis in APBs (ATSA) assay

ATSA assay was performed as previously described (Zhang et al., 2019) with slight modification. To synchronize cells in the G2 phase, cells were treated with 2.5 mM thymidine for 20 h, released into a fresh medium for 6 h, and then treated with 15 μM RO-3306 (CDK1i) for 16 h before fixation. The synchronization was confirmed by flow cytometry analysis of cells additionally labeled with 10 μM EdU for 30 min. For visualization of DNA synthesis in APBs, cells were incubated with 20 μM EdU for 30 min before fixation. The fixed cells were subjected to immunostaining for PML, EdU-click reaction, and followed telomere FISH with TelC-Cy3 probe.

### Mitotic DNA synthesis (MiDAS) assay

MiDAS assay was performed as previously described (Wu et al., 2020) with slight modification. After transfection, cells were treated with 0.4 μM aphidicolin for 16 h and 15 μM of RO-3306 (CDK1i) for 6 h before colcemid treatment. After washing with 1X PBS, cells were incubated in fresh culture media (to release cells from G2 arrest) containing 1X colcemid (to arrest cells at the metaphase) and 20 μM EdU (to label mitotic DNA synthesis) for 1 h before fixation. After trypsinization, cells were slowly mixed with pre-warmed 75 mM KCl in a drop-wise manner and then incubated at 37°C for 12 min. Cells were then fixed using a cold fixative solution (methanol/acetic acid 3:1). After fixation, cells in suspension were spread onto a clean slide and exposed to 90°C steam for 30 s. Slides were dried at room temperature for 1 h. Cells were then subjected to EdU-click reaction and followed telomere FISH with TelC-Cy3 probe.

### EdU-click chemistry for ATSA, MiDAS, and cell cycle analysis

Cells were labeled with 10∼20 μM EdU according to conditions in each assay. After EdU labeling, a click reaction was performed using the Click-iT® EdU Imaging kit (Thermo Fisher Scientific), according to the manufacturer’s instructions. For EdU-click reaction for MiDAS assay, cells were incubated with pre-extraction buffer (descripted in immunostaining section) for 10 min at 4°C. Cells were then rinsed with 1X cold-PBS, fixed with 4% PFA for 15 min, and permeabilized with 0.5% Triton X -100 for 5 min before click-reaction. For cell cycle analysis, cells were labeled with 10 μM EdU for 30 min before harvesting and processed using the Click-iT^®^ EdU flow cytometry assay kit (C10643) (Thermo Fisher). In brief, cells were fixed, permeabilized, and subjected to click reaction. Cells were then incubated in PBS with 0.1 mg/mL RNase A for 1 h at 37°C and DNA was stained with 0.05 mg/mL propidium iodide. Flow cytometry was performed on a FACSVerse™ flow cytometer using the BD FACSuite™ software (BD Biosciences). Data analysis was performed by using the FlowJo software.

### Image acquisition and image analysis

For all immunostaining assays combined with telomere FISH (except PCNA) or EdU-Click reaction, including ATSA, MiDAS, SMAT assay, and CO-FISH assay, images were acquired using a BX53 fluorescence microscope (Olympus LS) with cellSens v.1.18 imaging software and analyzed with OlyVIA software (Olympus LS). For PCNA immunostaining with telomere FISH, confocal images were acquired using an LSM880 confocal microscope (Carl Zeiss) with a 40×/1.2 lens objective. Image acquisition and analysis were performed with Zen 2.6 (blue edition) (Carl Zeiss) software.

### Flow cytometry and FISH (Flow-FISH)

Flow-FISH assay to measure the average length of telomeres was performed as previously described (Baerlocher et al., 2006) with slight modification. After trypsinization, 5x10^6^ cells were collected and incubated with hybridization buffer (70% formamide, 20 mM Tris-HCl, pH 6.8, and 1% BSA in 2X SSC) at room temperature for 10 min. For probe denaturation, 0.05 μg/ml PNA probe (TelC-FITC) was added to the hybridization buffer and denatured at 82°C for 5 min, and rapidly cooled in ice for 2 min. After incubation with a denatured probe for 2 h at room temperature in dark, cells were washed with washing buffer (70% formamide, 10 mM Tris-HCl, pH 6.8, 0.1% BSA in TDW, and 0.1% Tween-20 in 2X SSC) twice. After centrifugation, pellets were subjected to propidium iodide staining (0.1% BSA, 10 mg/ml RNase A, 0.1 mg/ml propidium iodide) for overnight at 4°C in dark. Flow cytometry was performed on a FACSVerse™ flow cytometer using the BD FACSuite™ software (BD Biosciences). Each experiment began with calibration beads with Quantum^TM^ FITC-5 MESF kits (#555p, Bangs Laboratories). Quantification and analysis were performed by using the QuickCal® v.3.0. software (Bangs Laboratories).

### C-circle amplification assay

The C-circle assay was performed as previously described (Henson et al., 2009) with slight modification. To isolate genomic DNA, cell pellets were resuspended with QCP lysis buffer [50 mM KCl, 10 mM Tris-HCl, pH 8.5, 2 mM MgCl_2_, 0.5% IGEPAL CA-630 (I8896, Sigma), 0.5% Tween-20 in DEPC-treated TDW with protease (#19155, QIAGEN, 0.053 AU/ml], and incubated with a shake at 1400 rpm at 56°C for 1 h. To inactivate the protease activity, lysed cells were subjected to heat treatment at 70°C for 20 min. For C-circle amplification, 10 μl of 0.1∼1 μg genomic DNA was mixed with 10 μl of 2X C-circle master mix [0.2 mg/ml BSA (NEB), 0.2% Tween-20, 2 mM dATP, 2 mM dGTP, 2 mM dTTP, 8 mM DTT, 2X ϕ29 buffer in DEPC-treated TDW] supplemented with ϕ29 polymerase (EP0091, Thermo Scientific) or DEPC-treated TDW. C-circle amplification was performed at 30°C for 8 h, followed by ϕ29 polymerase heat inactivation at 65°C for 20 min and cooling at 4°C using PCR thermocycler (Eppendorf). Amplified C-circle DNA was diluted with 2X SSC buffer and loaded onto a dot-blot apparatus bedded with Hybond N+ membrane (RPN2020B, Amersharm). Bounded DNA was crosslinked to the membrane by exposing the membrane to 1300 J/m^2^ UV (254 nm) using a UV crosslinker (Stratagene). The membrane was pre-hybridized with DIG Easy Hyb solution (Sigma-Aldrich) at 42°C for 1 h. Digoxigenin-labeled oligonucleotide probes (CCCTAA)_4_ were added to the DIG Easy Hyb solution at a final concentration of 100 ng/mL and incubated with gentle agitation at 42°C overnight. The membrane was sequentially washed with 0.1% SDS in 2X SSC at room temperature and 0.1% SDS in 0.2X SSC at 50°C for 20 min each with gentle agitation. The membrane was then incubated with HRP-conjugated anti-Digoxigenin-antibody for 6 h. After washing with TBST buffer, the signal was detected using enhanced a chemiluminescence reagent (Thermo Fisher Scientific) with an automated imaging system (ChemiDoc™; Bio-Rad Laboratories).

## Statistical analysis

Prism 9 (GraphPad Software) was used to generate graphs and analyze data. Error bars represent the standard deviation (s.d.) of the mean (mean ± s.d.). For statistics, we used one-way ANOVA, along with a post-hoc Tukey test or two-tailed unpaired Student’s *t*-test, as descripted in the figure legends; **** p < 0.0001, *** p < 0.001, ** p<0.01, * p < 0.05 and n.s: not significant. Statistical parameters are described in the figures.

---

**Extended Data Fig. 1.**
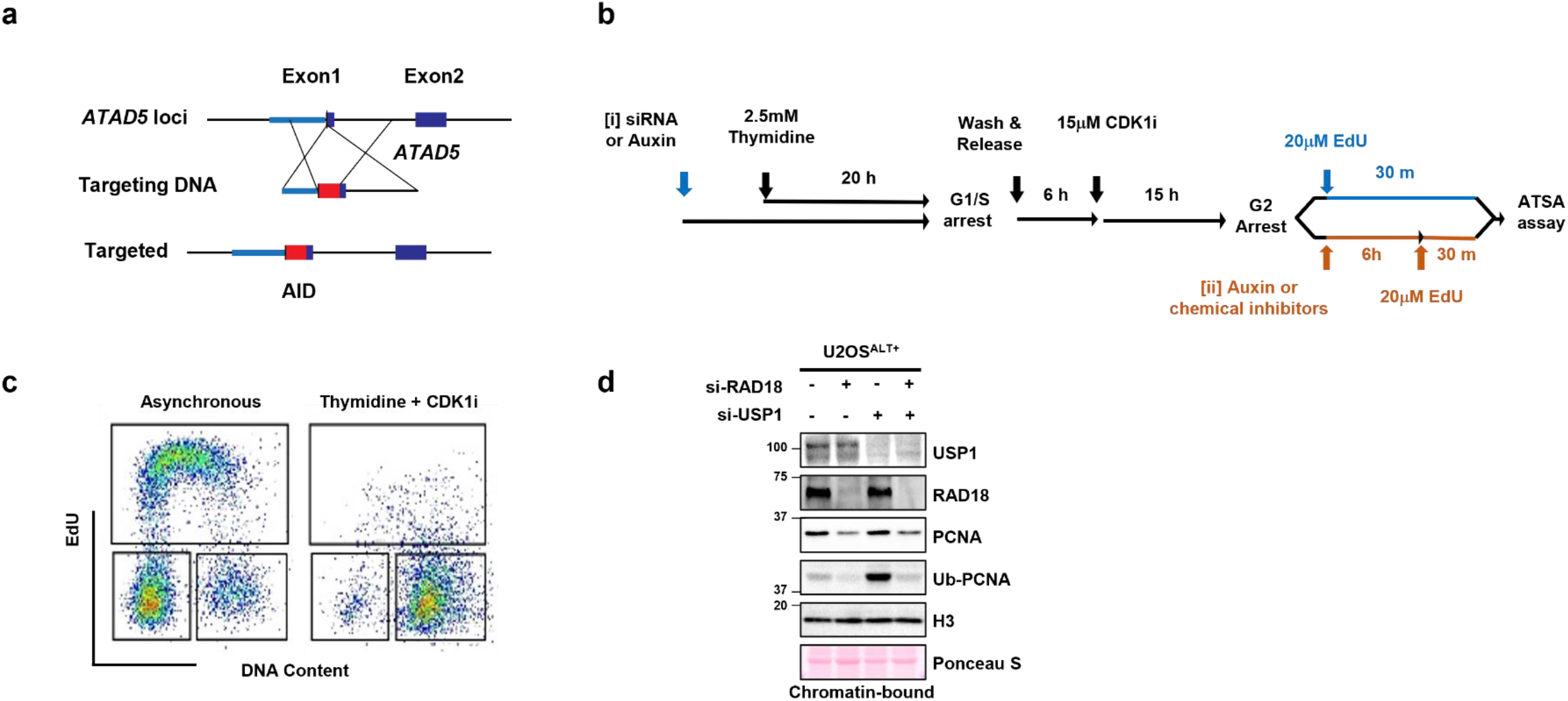
The experimental scheme for an ATSA assay and G2 synchronization, related to Fig. 1. (a) Genomic structure of the wild type (top) and the modified *ATAD5* locus (bottom) in U2OS- or HeLa-ATAD5^AID^ cell lines. (b) The experimental scheme for the G2 synchronization and an ATSA assay. (c) Representative cell cycle profile showing G2 synchronization. (d) After siRNA transfection for 48 h, U2OS cells were subjected to immunoblotting with chromatin-bound protein extracts.

**Extended Data Fig. 2.**
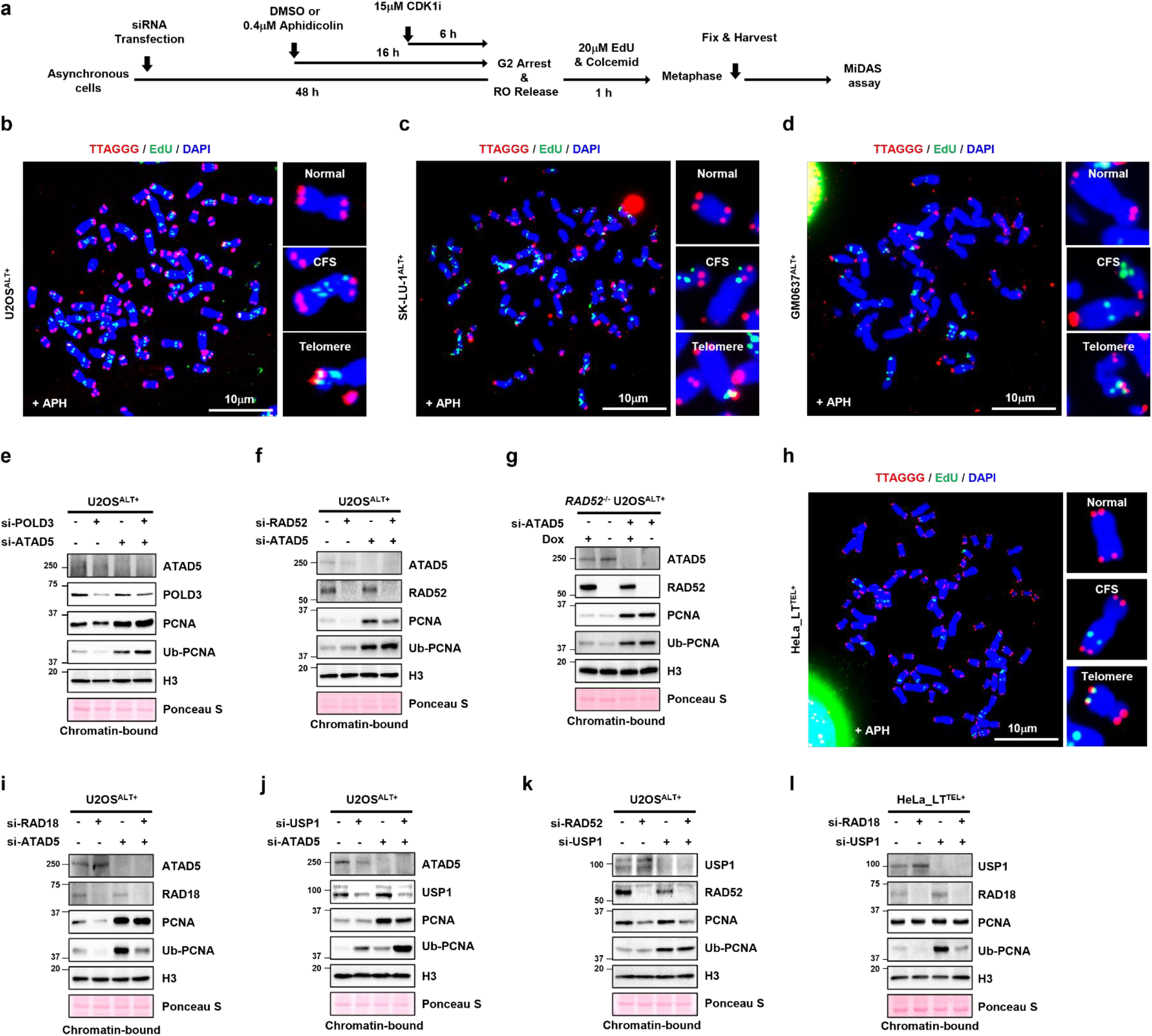
The experimental scheme for the MiDAS assay and biochemical characterization of the cells corresponding to each experiment, related to Fig. 2. (a) The experimental scheme for the MiDAS assay. (b-d, h) Representative images of MiDAS after APH treatment in corresponding cells. (e-g, i-l) After siRNA transfection, cells were subjected to immunoblotting with chromatin-bound protein extracts. (g) RAD52 expression was induced by doxycycline (DOX) for 48 h. Cell lines: U2OS (b, e, f, i-k); *RAD52*^−/−^ U2OS (g); SK-LU-1 (c); GM0637 (d); HeLa_LT (h, l).

**Extended Data Fig. 3.**
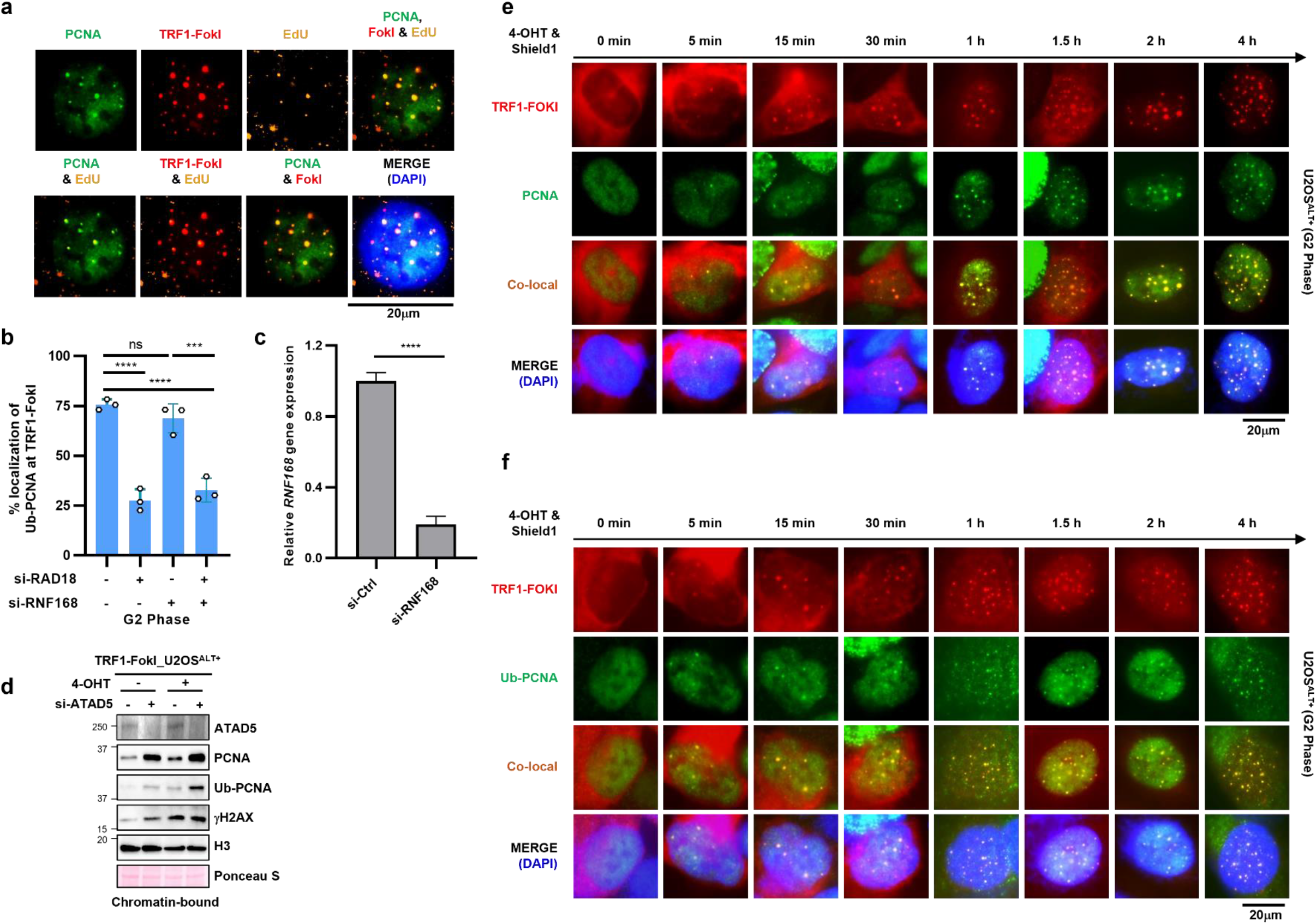
Ub-PCNA localization at TRF1-FokI foci is independent of RNF168 in ALT+ cancer cells, related to Fig. 3. (a, b, d, e, f) After siRNA transfection, U2OS-mCherry-TRF1-FokI cells were enriched in the G2 phase, Cells were then treated with tamoxifen and Shield1 for 2 h (a, b, d) or indicated times (e, f) before fixation for immunostaining. (a) Representative immunostaining images of PCNA, EdU, and TRF1-FokI co-localization after telomeric break-induction by TRF1-FokI. (b) The percent localization of Ub-PCNA at TRF1-FokI foci was quantified. (c) The knockdown efficiency of *RNF168* siRNA was quantified by RT-qPCR. (d) Chromatin-bound protein extracts were prepared for immunoblotting. (e, f) Representative images from cells transfected with *ATAD5* siRNA. Data in all graphs represent the mean ± s.d. of at least three independent experiments. Statistical analysis: One-way ANOVA (b); two-tailed unpaired Student’s *t*-test (c).

**Extended Data Fig. 4.**
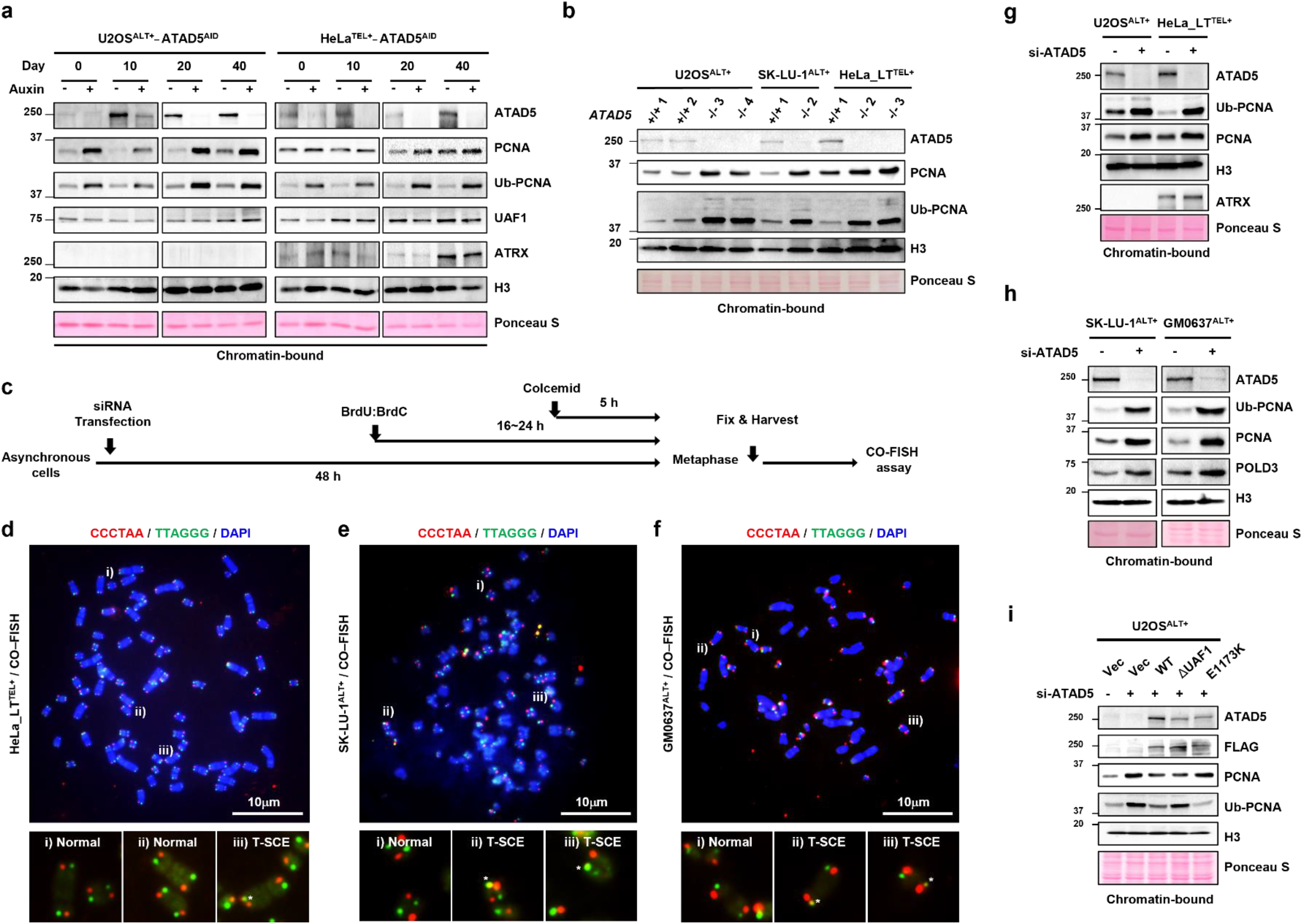
Biochemical characterization of *ATAD5* knockout and ATAD5-depleted cells, related to Fig. 4. (a) U2OS- or HeLa-ATAD5^AID^ cells were treated with auxin for the indicated days, and chromatin-bound protein extracts were prepared for immunoblotting. (b) *ATAD5*^+/+^ and *ATAD5^−/−^* clones of each cell line were subjected to immunoblotting with chromatin-bound protein extracts. (c) The experimental scheme for a CO-FISH assay to measure T-SCE. (d-f) Representative images of CO-FISH. (g, h) After siRNA transfection, indicated cells were subjected to immunoblotting with chromatin-bound protein extracts. (i) After transfection with a combination of *ATAD5* siRNA and wild type (WT), UAF1 interaction-defective (ΔUAF1) or PCNA unloading-defective (E1173K) ATAD5 cDNA, U2OS cells were subjected to immunoblotting with chromatin-bound protein extracts.

**Extended Data Fig. 5.**
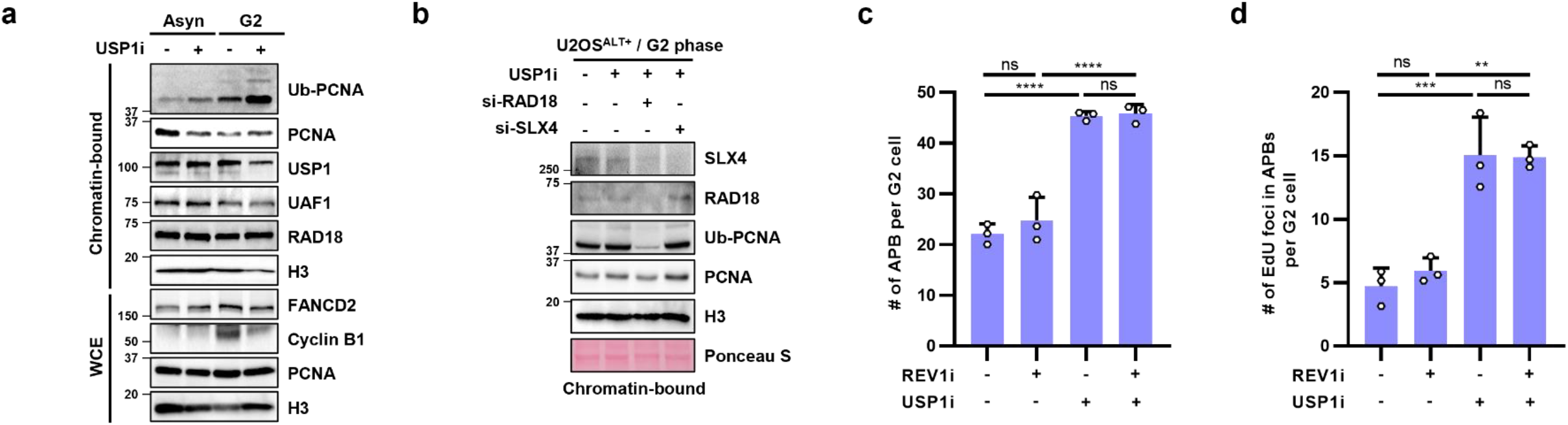
USP1 inhibitor increases monoUb-, and polyUb-PCNA in G2 phase of ALT+ cancer cells, related to Fig. 5. (a) U2OS cells asynchronous (Asyn) or synchronized in the G2 phase (G2) were treated with USP1i for 6 h and chromatin-bound protein extracts or whole cell extracts (WCE) were prepared. (b) U2OS cells were transfected with siRNAs as indicated, synchronized in the G2 phase, treated with USP1i for 6 h, and chromatin-bound protein extracts were prepared. (c-d) U2OS cells were synchronized in the G2 phase, treated with USP1i and/or REV1i for 6 h, and subjected to an ATSA assay. Data in all graphs represent the mean ± s.d. of at least three independent experiments. Statistical analysis: One-way ANOVA (c, d).

**Extended Data Fig. 6.**
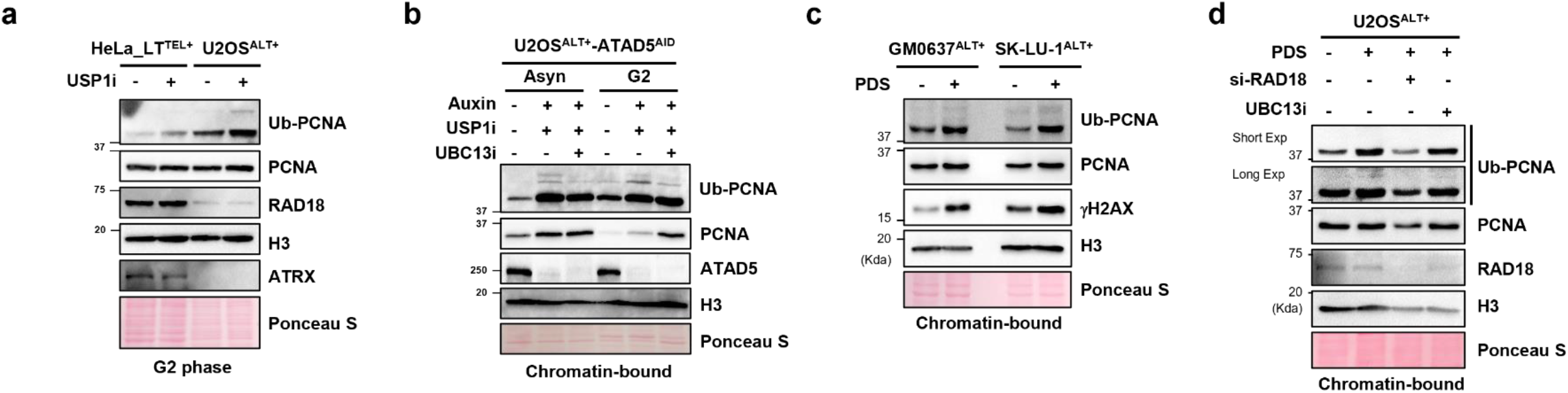
G4 stabilizer increases monoUb- and polyUb-PCNA in ALT+ cancer cells, related to Fig. 6. (a) HeLa_LT or U2OS cells synchronized in the G2 phase were treated with USP1i for 6 h, and whole cell extracts were prepared for immunoblotting. (b) Asynchronous (Asyn) or G2-synchronized (G2) U2OS-ATAD5^AID^ cells were treated with auxin, USP1i, and UBC13i for 6 h, and chromatin-bound proteins were extracted. (c) GM0637 and SK-LU-1 cells were treated with pyridostatin (PDS) for 24 h and subjected to immunoblotting with chromatin-bound protein extracts. (d) After siRNA transfection, U2OS cells were treated with pyridostatin (PDS) for 24 h and UBC13i for 6 h, respectively before extraction of chromatin-bound proteins.

**Extended Data Fig. 7.**
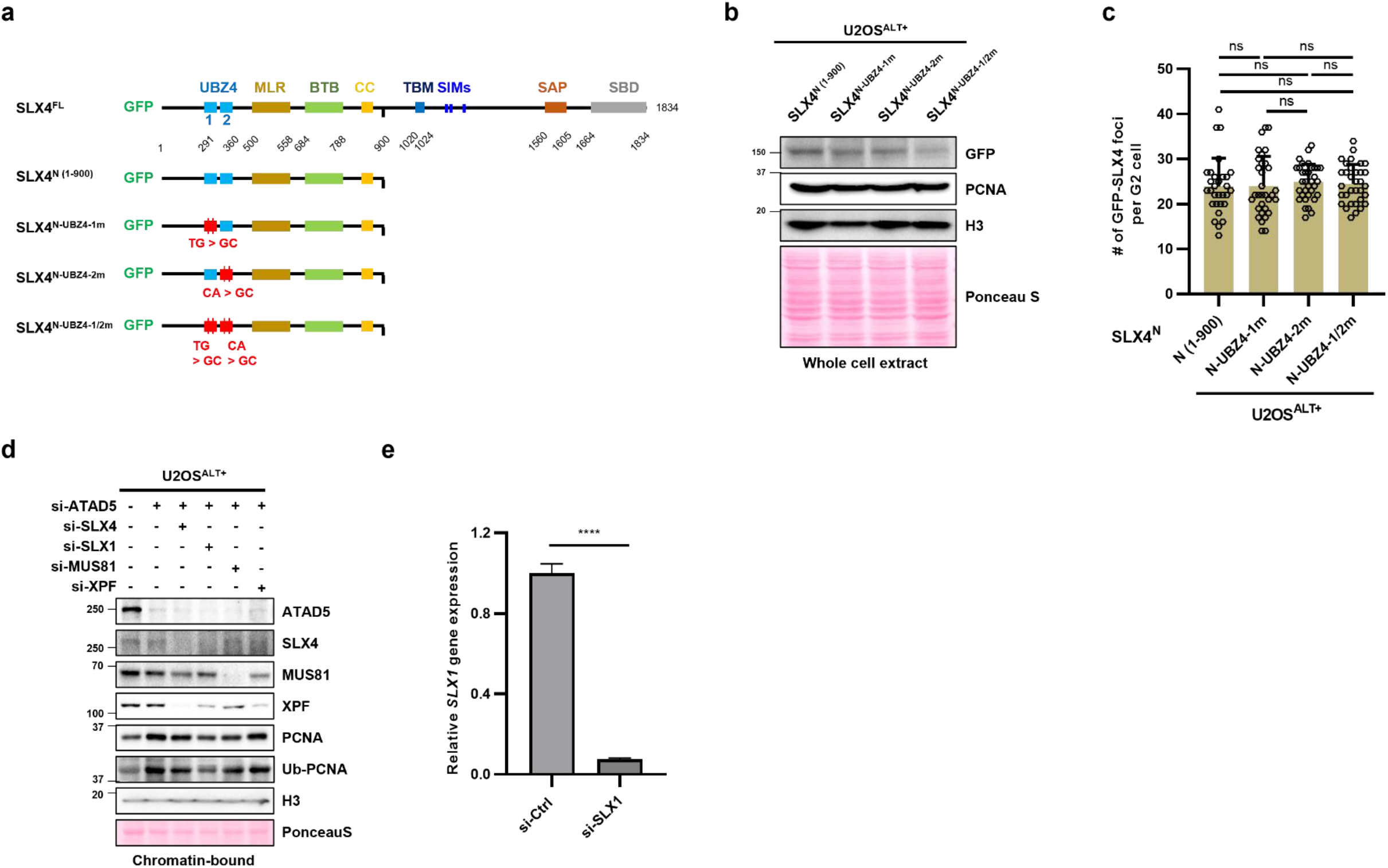
Characterization of U2OS cells expressing GFP-tagged SLX4 N-terminal fragment with mutations in UBZ4 domain, related to Fig. 6. (a) Schematic illustration of GFP-tagged full-length (FL) and N-terminal fragment (N) of SLX4 proteins and its functional domains (UBZ4: ubiquitin binding zinc fingers 4; MLR: Mus312-ME19-interaction-like region; BTB: *broad*-*complex*, *tramtrack, and bric*-à-*brac*; CC: coiled-coil; TBM: TRF2-binding motif; SIMs: SUMO-interacting motifs; SAP: SAF-A/B, Acinus, and PIAS; SBD: SLX1-binding domain). Mutations in the UBZ4 domain are highlighted in red. (b, c) U2OS cells expressing GFP-tagged wild type or mutant SLX4^N^ and subjected to immunoblotting (b) or synchronized in the G2 phase for immunostaining (c). (c) The number of GFP-SLX4 foci was quantified. (d) After transfection in U2OS cells, chromatin-bound protein extracts were prepared for immunoblotting. (e) The knockdown efficiency of *SLX1* siRNA was quantified by RT-qPCR analysis. Data in all graphs represent the mean ± s.d. of at least three independent experiments. Statistical analysis: two-tailed unpaired Student’s *t*-test (c, e).

**Extended Data Fig. 8.**
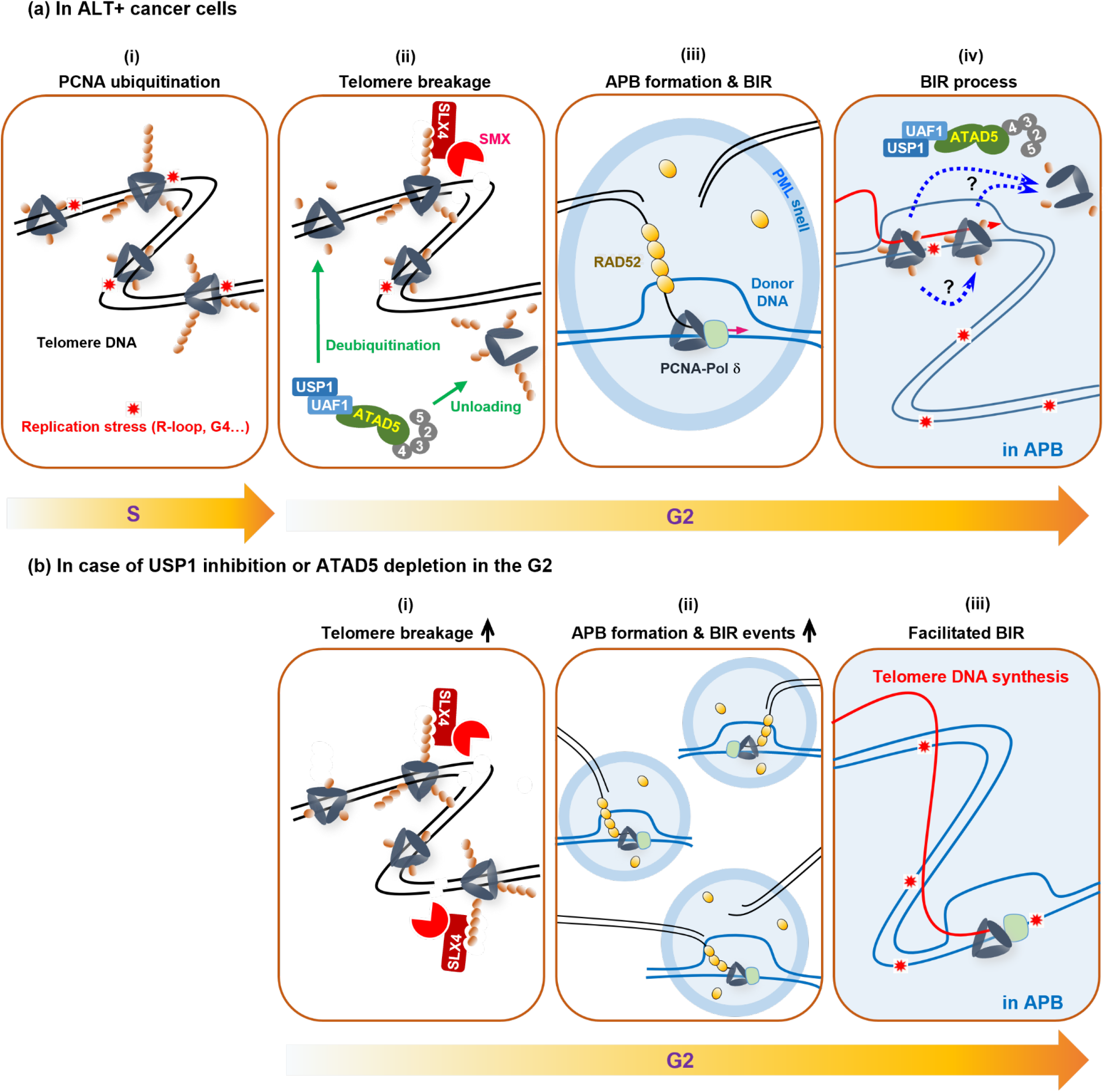
Graphical model of polyUb-PCNA-induced generation of telomere breakage and Ub-PCNA-induced promotion of the BIR processes in ALT+ cancer cells. [a, (i-ii)] Telomeres in ALT+ cells suffer more frequent replication stress than those in TEL+ cells. MonoUb- and polyUb-PCNA, formed under telomeric replication stress, remain on chromatin at a significant level until the G2 phase due to an unknown mechanism that restricts unloading and deubiquitination of Ub-PCNA. Although it is not clear whether monoUb- and polyUb-PCNA in the G2 phase have completed their roles in TLS- and template switch-mediated damage bypass, respectively, the K63-linked-polyubiquitin chain of PCNA additionally acts as a binding site for SLX4 via the UBZ4 domain and consequently acts as a working platform for the SLX4-associated SMX tri-nuclease complex. All three endonucleases comprising the SMX tri-nuclease complex (SLX1, MUS81, and XPF) cooperatively generate telomere breakage. [a, (iii)] Broken telomeric DNA ends induce APB formation with a help of BLM helicase and PML, and then RAD52- and POLD3-dependent BIR is initiated in APBs. [a, (ii)] Telomeric breakage by polyUb-PCNA-SLX4 is inhibited by balanced deubiquitination and/or unloading of polyUb-PCNA by the USP1-UAF1-ATAD5-RLC. [b, (i-ii)] When this counterbalance is disrupted by acute ATAD5 depletion or USP1 inhibition, even in the G2 phase, the excess polyUb-PCNA increases the abundance of the SLX4 and SMX tri-nuclease complex at telomeres, which leads to sequential increases in telomeric breakage, APB formation, ALT phenotypes, and telomere length. [A, (iv)] During the BIR process, PCNA-polymerase δ encounters replication stress, and, consequently, PCNA is ubiquitinated. Although whether monoUb-PCNA or polyUb-PCNA is present and the role of Ub-PCNA during BIR is unclear, our results suggest that Ub-PCNA facilitates the BIR process. [B, (iii)] Consistently, Ub-PCNA left unprocessed due to depletion of ATAD5 or USP1 increased TRF1-FokI-induced telomeric DNA synthesis and telomeric MiDAS.

## Notes

### Competing Interest Statement

The authors have declared no competing interest.

